# Compressive stress couples calcium signalling to translational control and proteostasis

**DOI:** 10.64898/2026.10.07.757237

**Authors:** Giuseppe Fragale, Benjamin J. Hershey, Yuwei Li, Giorgia Parodi, Galina V. Beznoussenko, Elisa Allievi, Arta Ajazi, Nicol Berti, Wanxin Zheng, Zeno Lavagnino, Vittoria Matafora, Federica Pisati, Nicola Manfrini, Stefano Biffo, Ligia Martinez, Andromachi Karakatsani, Martina Guadagno, Andreas E. Moor, Dario Parazzoli, Marco Cosentino-Lagomarsino, Davide Mazza, Angela Bachi, Paolo Maiuri, Kristina M. Havas

**Affiliations:** Istituto di Genetica Molecolare, Consiglio Nazionale delle Ricerche (IGM-CNR), Via Abbiategrasso 207, Pavia, Italy; IFOM ETS-The AIRC Institute of Molecular Oncology, Via Adamello 16, 20139, Milan, Italy; ETH Zurich, Department of Biosystems Science and Engineering, Klingelbergstrasse 48, Basel, Switzerland; Cogentech SRL Benefit Corporation, Milan, Italy 20139; INGM, Istituto Nazionale Genetica Molecolare Romeo ed Enrica Invernizzi, Milan, Italy; Department of Biosciences, University of Milan, Milan, Italy; Dipartimento di Fisica, Università degli Studi di Milano, Milan, Italy; INFN – Istituto Nazionale Fisica Nucleare sezione di Milano, Milan, Italy; Università Vita-Salute San Raffaele, Milan, Italy; Experimental Imaging Center, IRCCS Ospedale San Raffaele, Milan, Italy; Department of Molecular Medicine and Medical Biotechnology, Università degli Studi di Napoli Federico II, Naples, Italy

## Abstract

Compressive stress is a pervasive feature of solid tumour growth, and mechanical loading is known to activate calcium signalling through the mechanosensitive channel Piezo1. How this signal is converted into changes in cellular biosynthetic capacity, however, remains unclear. Here we show that compressive stress engages a Piezo1-dependent calcium checkpoint that reduces translational output with increasing mechanical load, independently of mTORC1 and the integrated stress response. Compression triggers Piezo1-dependent release of endoplasmic reticulum Ca²⁺, activating eEF2 kinase to slow translational elongation. This same Ca²⁺ signal drives mitochondrial remodelling and reactive oxygen species production, which reinforce translational repression through redox-dependent oxidation of the translation initiation factor eIF2β. In parallel, Ca²⁺-dependent AMPK activation triggers autophagic ribosome degradation, alleviating compression-induced macromolecular crowding and restoring cytoplasmic rheology despite sustained mechanical load. These mechanisms act at distinct levels of translational control: initiation, elongation and ribosome abundance; yet converge to limit biosynthetic output under compression. Consistent with a role for this pathway in tumour adaptation to mechanical constraint, eEF2K expression correlates with poor overall survival in a public PDAC patient dataset. Together, these findings define a mechano-metabolic checkpoint through which cells actively tune biosynthetic capacity to their physical environment.

## Introduction

Cells inhabit mechanically dynamic environments where forces such as tension, shear stress, and compression continuously shape their behaviour, metabolism, and fate (*1–7*). While tensile and shear forces have been extensively characterized as drivers of mechanotransduction, compressive stress remains a comparatively underexplored mode of mechanical signalling, particularly regarding its influence on intracellular biochemistry and biomass regulation. Despite this, compression is a pervasive physical constraint that arises in both physiological and pathological contexts as cells proliferate, migrate, or remodel within confined environments. During gut morphogenesis, for instance, circumferential contraction of smooth muscle layers generates compressive forces on the expanding endoderm and mesenchyme, driving villus formation through coordinated buckling and folding (*8*, *9*). In solid tumours, growth-induced stresses can reach several kilopascals, profoundly influencing tumour architecture, progression and therapeutic response (*10–14*). While moderate compression serves as a key regulator of cell behaviour by shaping developmental trajectories and tissue organization in both physiological and pathological contexts, excessive levels of compressive stress has been shown to trigger apoptotic and necrotic programs(*15*, *16*). Indeed, most nascent tumours fail to advance to overt malignancy, and failure to adapt to growth-induced mechanical stress may in part explain this restraint. The critical question, therefore, is how the tumours that do progress manage to overcome these biophysical barriers to growth.

Mechanical compression alters cell shape, volume, and intracellular crowding, all of which can affect the diffusion of macromolecules, the organization of organelles, and the efficiency of enzymatic reactions(*17*). Because most biochemical reactions depend on molecular mobility and crowding, these physical changes threaten core homeostatic processes including protein synthesis and degradation. Protein synthesis is one of the most energetically demanding cellular activities and principal driver of biomass accumulation, its associated machinery also contributes to macromolecular crowding(*18–20*). Translational regulation allows cells to rapidly adjust protein synthesis rates in response to fluctuations in nutrients, energy, and stress, thereby controlling growth and survival (*21*). Because of this, it represents a prime regulatory node for adapting to mechanical challenges. While nutrient deprivation and energy limitation are known to regulate translation via mTORC1 and the integrated stress response, how compressive stress is sensed and transduced into biochemical adaptation remains poorly understood.

Here, we investigate how cells adapt to compressive stress that reduces volume and increases macromolecular crowding. We show that mechanical compression elicits calcium-dependent signalling that translates physical deformation into coordinated biochemical responses regulating protein synthesis and proteostasis. Acute compression rapidly tunes protein synthesis, through activation of the calcium/calmodulin-dependent kinase eEF2K, independently of mTORC1 and the integrated stress response. This acute response is reinforced by Piezo1-dependent mitochondrial remodelling and reactive oxygen species production, which sustain translational inhibition through oxidation of the redox-sensitive component of the ternary complex, eIF2β. In parallel, compression induced Ca^2+^ flux promotes the re-establishment of cytoplasmic rheological homeostasis via AMPK-dependent autophagic clearance of ribosomes. Together, these findings reveal that cells exposed to compressive stresses engage a coordinated mechanical checkpoint coupling calcium signalling, redox metabolism, and translational control to preserve biomass homeostasis in physically constrained environments.

## Results

### Compressive stress increases macromolecular crowding and suppresses translation

To determine how compressive stresses affect cell volume and macromolecular crowding we subjected colorectal cancer cell line HCT116 to 1D compression using elastic overlays constructed from polyacrylamide gels at stiffnesses ranging from 2-30 kPa (Fig. 1A)(*22*, *23*). HCT116, widely used in therapeutic research, provide a tractable model of cells adapted to growth under chronic compressive stress, a defining mechanical feature of solid tumours(*24*). Nuclear volume, used as a proxy for cell volume, showed a persistent stiffness-dependent reduction (Fig. 1B), suggesting that this compression mode is also volumetric. The average nuclear volume decreased by 4% in 2kPa, 15% in 10 kPa up to 43% in 30 kPa compressed cells (*V*_cntrl_=1288±352µm^3^, *V*_2kPa_=1,233±328µm^3,^ *V*_10kPa_=1,092±270µm^3^ and *V*_30kPa_=737±266µm^3^). It has been previously reported that volume loss upon cell confinement primarily reflects water efflux (*17*), therefore we predicted a concomitant increase in macromolecular crowding upon compression.

**Figure 1.**
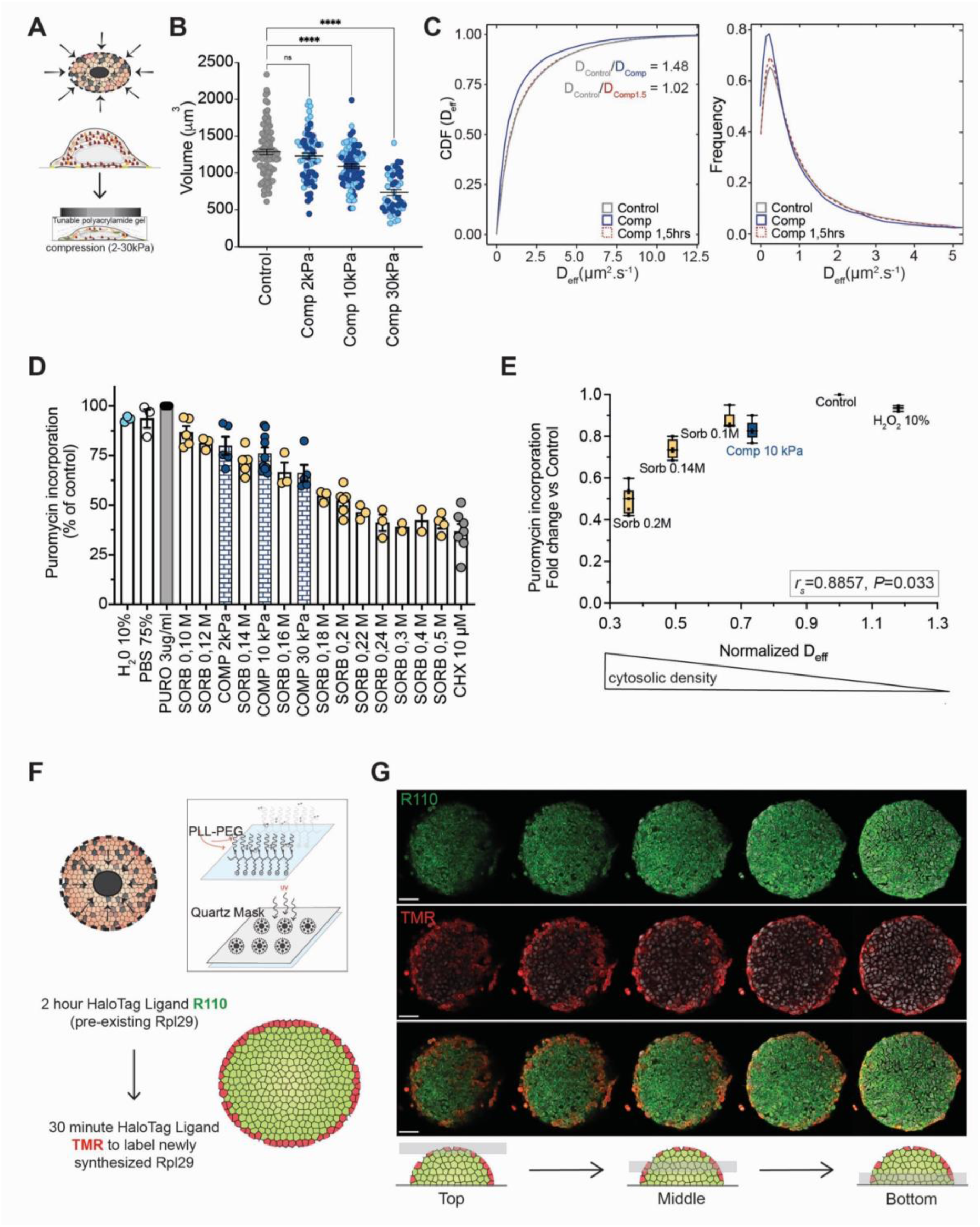
Mechanical tuning of cell biosynthetic processes. **A.** Schematic illustrating externally applied and growth-induced solid stresses in solid tumours, and the tuneable polyacrylamide system used to apply external 1D solid stress to cells. **B.** Nuclear volumes measured in live cells under control, 2 kPa, 10 kPa, and 30 kPa compression conditions. Light blue circles represent single-cell nuclear volumes measured during the first 30 minutes following compression onset; dark blue circles represent those measured 30–60 minutes post-compression. Data are presented as mean ± SEM for data acquired in n=2 independent experiments. **C.** Cumulative distribution function of effective diffusion coefficients (*D*ₑ_ff_) for 40 nM GEMs under control (grey), compressed 0–30 minute (blue), and compressed 1.5 h (red) conditions. Distributions of *D*ₑ_ff_ values for individual GEMs are shown for the same conditions. Data represents pooled data from n=2 independent experiments. **D.** Puromycin incorporation shown as percentage relative to control for the indicated conditions. Points represent independent experiments; data are presented as mean ± SEM, for Sorbitol 0.3M and Sorbitol 0.4M n=2, for all other conditions n≥3 independent experiments. **E.** Plot of normalized median D_eff_ measured with 40 nm GEMs(n=2) versus normalized puromycin incorporation (n≥3) for the indicated conditions of osmotic shock and compression**. F.** Schematic of micropatterns and HCT116 RPL29_HALO line used to measure translation under growth induced compression. **G.** Representative images of HCT116 RPL29_HALO cells grown on 250 µm disks, with R110 labeling of pre-existing RPL29 followed by a 30-minute TMR pulse to label newly synthesized protein. Scale bar: 50 µm. One way ANOVA with Sidak’s multiple comparisons test(B), Spearman r Correlation analysis (E). *P values: ∗∗∗∗ P < 0.0001*.

To test the micro-rheological properties of the cytoplasm of compressed cells we used 40 nm genetically encoded multimeric nanoparticles (GEMs)(*25*). Single particle tracking of the GEMs was done at a 10ms framerate and the effective coefficients of diffusion (D_eff_) were extracted for 100ms time scales in both control and compressed conditions. We observed a significant decrease in the diffusivity of the GEMs measured within the first 30 minutes following the application of 10 kPa compression (D_cntrl_=0.888 µm^2^s^-1^, D_10kPa/0min_=0.602 µm^2^s^-1^_;_ Fig. 1C). We then asked whether the observed increase in macromolecular crowding remained constant in cells subjected to continuous compressive stress over a period of 90 minutes. Despite the sustained volume reduction, GEM diffusivity measured at 90 minutes returned to baseline values (D_10kPa/90min_= 0.867 µm^2^s^-1^; Fig. 1C), suggesting an active mechanism was driving the re-establishment of rheological properties of the compressed cells.

Proteins are the largest contributors to a cells dry mass, accounting for nearly 60% of the dry mass of a mammalian cell (*26*). Eukaryotic protein levels are primarily regulated at the level of mRNA translation, an energetically costly process that is tightly controlled to adapt to cellular responses to environmental stress. Therefore, we asked whether mechanical compression impacted translation by measuring puromycin incorporation following acute application of both compressive and osmotic stresses. We observed a progressive reduction in puromycilated peptides with increasing osmotic stress. Similarly, compressive stress induced a stiffness-dependent decrease in puromycin incorporation across the three conditions tested (2kPa (–23±4.6%), 10kPa (–31±1.7%), 30kPa (–38±1.3%)) (Fig. 1D and Supplementary Fig. 1A). This effect was conserved among multiple cell lines, including HeLa and human retinal epithelial cells (hTERT RPE-1) (Supplementary Fig. 1B,C). In agreement with the reduction in translation, we observed a significant increase in P-Body number and size within the first 15 minutes following compression, although a similar trend was not observed for stress granules (Supplementary Fig. 1D,E). To assess whether the translational capacity correlated to cytoplasmic density we performed single particle tracking of GEMs under a range of osmotic stresses from 10% H_2_O to 0.5M Sorbitol and under acute 10kPa compression. We then plotted the normalized D_eff_ versus puromycin incorporation and observed a positive correlation between cytoplasmic density and translational capacity (*r_s_ =*0.886) (Fig. 1E). Taken together this data suggested that externally applied compressive stress results in volume driven increased macromolecular crowding and reduced translational output.

Compressive stress generated by tumour growth may therefore provide a physiological mechanism for regulating translational capacity (Fig. 1A)(*27*). To address whether growth-induced compression can tune translation in a spatially defined context, we chose to utilize micropatterning *in lieu* of organoid-based models to minimize confounding effects intrinsic to organoid cultures, such as limited nutrient diffusion that could affect translation. We seeded HCT116 cells engineered via CRISPR–Cas9 to express a HALO tag at the C-terminus of endogenous RPL29 (HCT116 Ribo-HALO) (*28*), onto 250 µm micropatterned disks, surrounded by a non-adhesive surface and allowed them to grow to confluency (Fig. 1F) (*29*). The use of micropatterns limited the growth area allowing for the accumulation of growth induced solid stress at the center of the pattern as quantified by a reduction in nuclear area (Supplementary Fig. 1F). HALO tagged RPL29 was used to measure translation capacity across the micropatterned disk. First, pre-existing RPL29 was irreversibly labeled with HaloTag ligand R110, followed by labeling of newly synthesized RPL29 with a second HaloTag ligand, TMR (Fig. 1G). We observed a distinct enrichment of the TMR signal in the cells at the periphery of the micropatterned disks, suggesting that growth induced pressure within the center of the disk limits translational output (Fig. 1G). By analyzing EdU incorporation we demonstrated that observed differences were not due to reduction in proliferation within the micropatterned disk (Supplementary Fig. 1). We concluded that compressive stress, whether externally applied, or arising from growth induced pressure, actively tunes translation rates.

### Acute translational repression is independent of mTORC1 and ISR

Motivated by mTOR’s role in translational control and its mechano-sensitivity, we asked whether mTOR activity was acutely modulated by mechanical stress (Fig 2A) (*30–33*). HCT116 cells were subjected to 10kPa compressive stress, and phosphorylation of mTORC1 targets p70S6Kinase (pT389) and 4EBP1 (pT37/46) as well as mTORC2 target Akt (pS473), was monitored over a three-hour time course. Consistent with previous reports, compressive stress induced a rapid and transient increase in Akt S473 phosphorylation within 10 minutes of compression, indicating an early mechano-activation of mTORC2(Fig. 2B) (*34*, *35*). In contrast, phosphorylation of the mTORC1 targets p70S6K and 4EBP1 was maintained during the first 60 minutes of compression, with significant dephosphorylation occurring only at later time points (≥60 minutes). These data indicate that mTORC1 activity is not acutely regulated by mechanical stress and that mechano-dependent mTORC1 inactivation is not an immediate early response. By comparison, hyperosmotic shock (0.25M Sorbitol) lead to rapid mTORC1 inactivation within 15 minutes (Supplementary Fig. 2A). Together, these results demonstrate that while mTOR complexes are mechanoresponsive, mTORC1 activity is not modulated on short time scales by compressive stress, and osmotic stress does not fully recapitulate compression-induced signaling.

**Figure 2.**
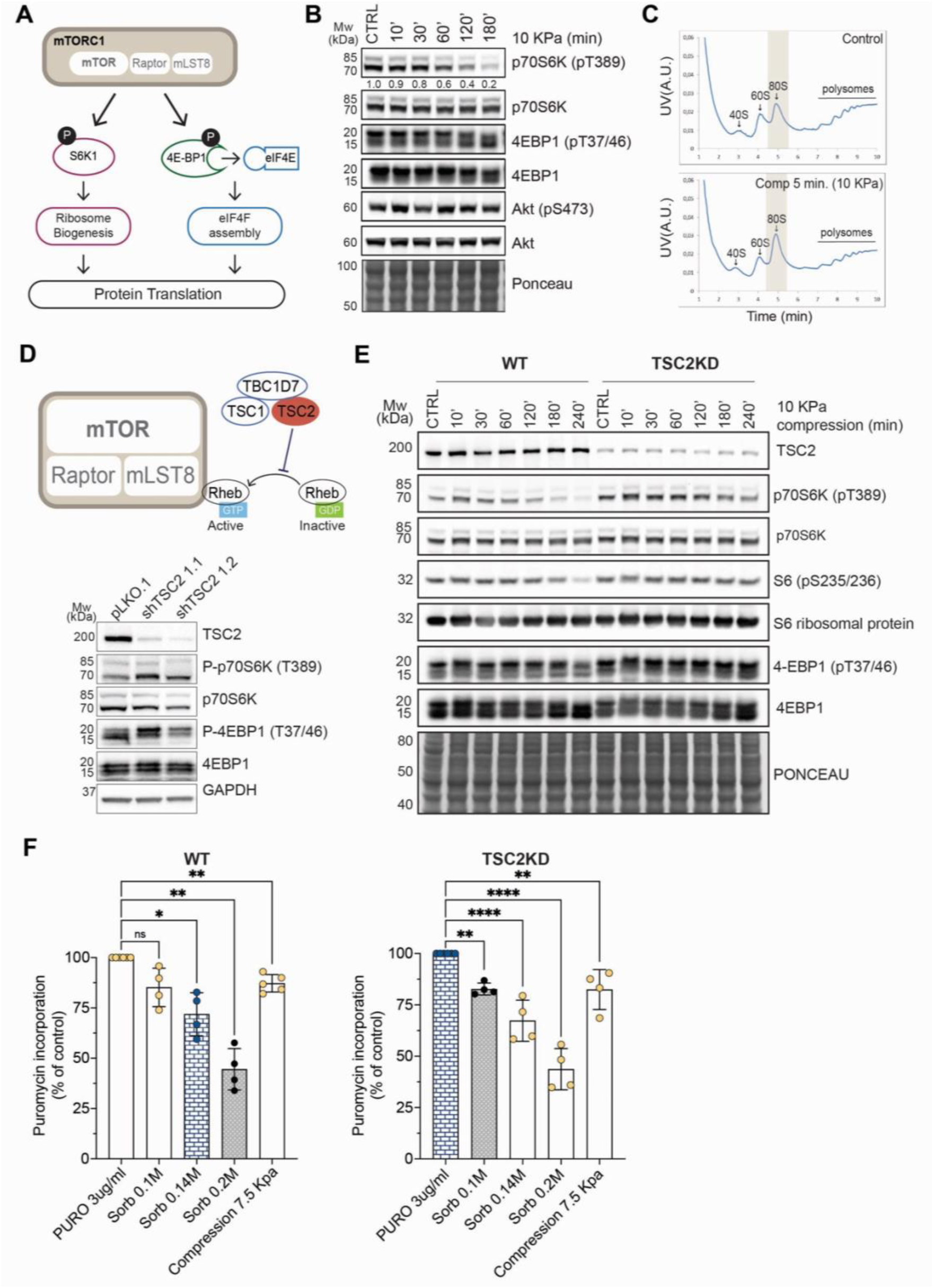
mTORC1 is dispensable for initial compression induced translation inhibition. **A.** Schematic of mTORC1 function in the regulation of translation over phosphorylation of S6K and 4E-BP1. **B.** Compression (10KPa) time course of mTORC1 substrates p70S6K(T389) and 4EBP1(T37/46) and mTORC2 substrate Akt (S473) phosphorylation following compression onset. Ponceau S staining serves as a loading control. Blots are representative of n=3 independent experiments. **C.** Polysome profiling of control and <5 minute compressed samples. Grey area highlights compression driven changes in 80S monosome levels. **D.** Illustration of TSC2’s role as a negative regulator of mTORC1 activity, and representative western showing heightened phosphorylation of mTORC1 targets p70S6K and 4EBP1 in TSC2kd cells. GAPDH serves as a loading control. **E.** Compression (10kPa) time course of mTORC1 substrates p70S6K(T389), S6(S235/236) and 4EBP1(T37/46) in control and TSC2kd cells. Ponceau S staining serves as a loading control. Blots are representative of n=3 independent experiments. **F.** Puromycin incorporation normalized to control in control and TSC2kd cells for the indicated conditions. Data points represent independent experiments, plotted as Mean ± SEM for n≥3 independent experiments. One-way ANOVA with Tukey’s multiple comparison test(F). *P* values: ∗ *P* < 0.05, ∗∗ *P* < 0.01, ∗∗∗ *P* < 0.001, ∗∗∗∗p *P* < 0.0001.

To test whether translational repression precedes mTORC1 inactivation, we performed polysome profiling on control and <5-minute 10 kPa compressed samples. Compression induced reduced polysome formation and accumulation of the 80S monosome as early alteration in translation dynamics, suggesting a reduced preceding mTORC1 inactivation (Fig. 2C).

To further demonstrate that acute mechano-repression of translation occurs independently of mTORC1, we knocked down Tuberous sclerosis complex 2 (TSC2), to constitutively activate mTORC1 (Fig. 2D). In TSC2-knockdown (kd) cells, mTORC1 targets showed heightened baseline phosphorylation that remained sustained after compression, whereas control (pLKO sh-Scrambled) cells exhibited a gradual loss of mTORC1 activity similar to wild-type cells (Fig. 2E). Notably, even TSC2kd cells showed a gradual reduction in the phosphorylation of mTORC1 targets, albeit at much later timepoints, suggesting a TSC2-independent mechanism for eventual silencing. Although mTORC1 signaling remained elevated in TSC2-depleted cells under compression, translational repression persisted under both 10kPa compression and hyperosmotic shock in TSC2kd cells, reinforcing the conclusion that mechano-induced translational repression occurs independently of mTORC1 (Fig. 2F).

We next examined the contribution of the integrated stress response (ISR) to early translational repression. The ISR is activated by diverse stresses and is executed through the phosphorylation of eIF2α leading to global reduction in protein synthesis, and induction of ATF4 translation (Supplementary Fig. 2B). We assessed the phosphorylation of eIF2α together with the activation state of several key kinases involved in the ISR pathway. We detected only a slight increase in eIF2α phosphorylation upon compression in both control and TSC2kd cells, with no significant activation of PERK or IRE1, and no early induction of ATF4 (Supplementary Fig. 2C). Treatment with 220 nM ISRIB, which antagonizes ISR by competing with phosphorylated eIF2α for eIF2B binding (Supplementary Fig. 2B and (*36*)), failed to rescue compression induced translational repression (Supplementary Fig. 2D).

Taken together, these results suggest that while mTORC1 and ISR may be modulated by compressive stress over longer timescales, both pathways are dispensable for the acute translational repression triggered by mechanical confinement.

### eEF2 Kinase is activated in response to compressive stress to tune translation

Having ruled out ISR and mTORC1 as the primary effectors of compression-induced translational inhibition, we performed an unbiased approach to identify phosphorylation events triggered by mechanical stress using phospho-proteomic analysis. We profiled control and 10kPa compressed cells. Sequential enrichment of phospho-peptides was carried out using Ti-IMAC and TiO_2_beads, followed by LC-MS/MS analysis (Fig. 3A). This approach identified 3,013 significantly differentially phosphorylated peptides, including 675 that showed a loss of phosphorylation and 2,338 that exhibited compression-induced phosphorylation. (Fig. 3B, Supplementary Fig. 3A, and Supplementary Dataset 1). GO enrichment analysis of proteins involved in translation revealed a differential phosphorylation state of proteins involved in several aspects of translational regulation, including ribosome biogenesis, mRNA stability and translation initiation and elongation (Supplementary Fig. 3B). Among these both eukaryotic elongation factor 2 (eEF2) and its regulatory kinase eEF2 kinase exhibited increased phosphorylation (Fig. 3B).

**Figure 3.**
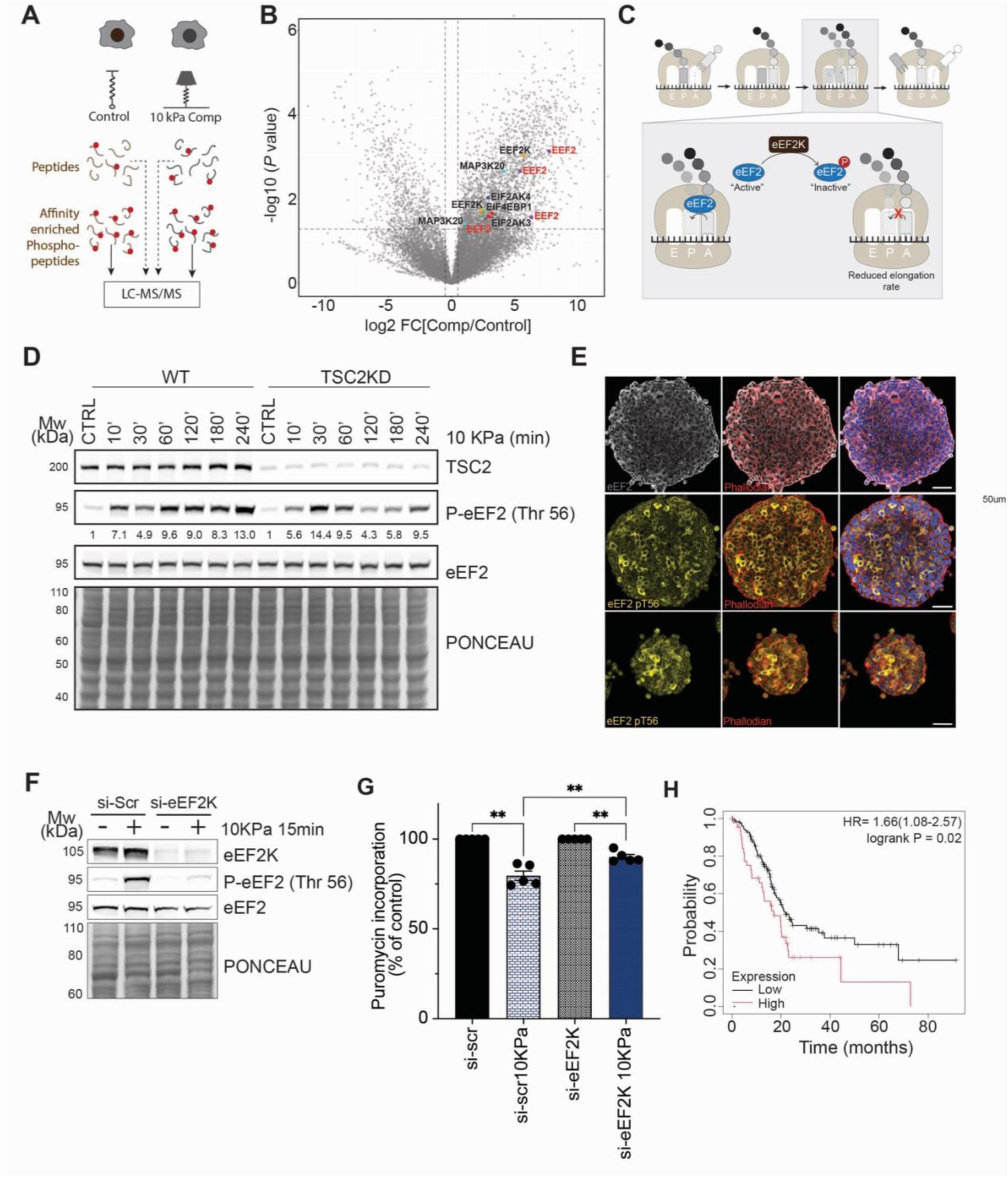
eEF2 kinase is a mechanosensitive regulator of protein biosynthesis. **A.** Schematic outlining the 2-step enrichment protocol used for the phospho-proteomic screen employed to identify compression induced alterations in the phospho-proteome. **B.** Volcano plot of significantly dysregulated phospho-peptides identified in the screen. Proteins implicated in translation are highlighted**. C.** Illustration of the role of eEF2 and its regulatory kinase eEF2 kinase in regulating the elongation step of protein translation. **D.** Western blot illustrating the temporal dynamics of eEF2 T56 phosphorylation upon 10kPa compression in control and TSC2kd cells. Ponceau S staining serves as a loading control. Blots are representative of n=3 independent experiments. **E.** Immunofluorescence HCT116 cells grown to confluence on 250 µm (top,middle) and 100 µM (bottom) disks for eEF2 PAN (grey) and eEF2 pT56(yellow) The scale bars represent 50 *μ*m. Representative of n=2 independent experiments. **F.** Western blot for eEF2 pT56 in control and si-eEF2Kinase cells under control and 10 kPa compressed conditions. Ponceau S staining serves as a loading control. Blots are representative of n=3 independent experiments. **G.** Quantified puromycin incorporation normalized to controls for control (si-scr) and si-eEF2Kinase in control and 10kPa compressed conditions. Data points represent independent experiments, plotted as Mean ± SEM for n=5 independent experiments. **H.** Kaplan Meier plot of overall survival for high versus low eEF2K expressing Pancreatic Ductal Adenocarcinoma patients (*38*). One-way ANOVA with Sidak’s multiple comparison test (G). *P* values: ∗p < 0.05

eEF2 is essential for the elongation phase of cytoplasmic protein translation, where it facilitates ribosomal translocation by shifting the peptidyl-tRNA from the A-site to the P-site, allowing entry of the next aminoacyl-tRNA. Its recruitment to the ribosome is negatively regulated through phosphorylation at threonine 56 by eEF2 kinase(*37*). By phosphorylating and inactivating eEF2, eEF2K slows down the elongation phase of translation (Fig. 3C). Given this regulatory role, mechano-activation of eEF2 kinase represents a strong candidate mechanism for modulating translation in response to mechanical stress prompting us to examine its regulation under compression. Given that compression induced translational inhibition was independent of mTORC1 we sought to validate this finding in both control and TSC2kd lines. Both lines were subjected to 10 kPa compression and eEF2 threonine 56 phosphorylation was monitored over a time course. We observed an induction of phosphorylation as early as 10 minutes following compression onset in both cell lines regardless of mTORC1 activation state, validating eEF2 phosphorylation following acute mechanical stress (Fig. 3D).

We then asked whether eEF2 phosphorylation could be in part responsible for the translational inhibition induced by growth induced pressure as observed in micropatterned cells at confluence. Immunofluorescence analysis of pThr56 eEF2 in micropatterns revealed an accumulation of eEF2 signal in centrally compressed regions mirroring the spatial translation repression observed in Fig. 1G (Fig. 3E). These findings suggest that eEF2 kinase is activated in response to mechanical compression.

To confirm its role as the upstream effector, we performed both pharmacological inhibition and siRNA-mediated knockdown of eEF2K. As expected, eEF2K-depleted cells exhibited significantly reduced phosphorylation of eEF2 at threonine 56 following compressive stress (Fig. 3F and Supplementary Fig. 3C,D). Finally, to directly link eEF2 kinase activation to the translational inhibition induced by compressive stress, we performed puromycin incorporation assays in control and eEF2K knockdown cells. Notably, eEF2K-depleted cells exhibited a significant rescue of the translation inhibition observed under compression in control cells, supporting a role for eEF2K as a mechano-sensitive kinase mediating translational repression in response to mechanical stress (Fig. 3G). However, because reduced elongation would be expected to increase ribosome occupancy, the polysome collapse observed following compression (Fig. 2C) suggests that eEF2K-dependent elongation inhibition does not fully account for the translational response.

Intriguingly, analysis of publicly available patient datasets revealed that increased eEF2K expression correlates with poorer overall survival in pancreatic ductal adenocarcinoma (PDAC), one of the solid tumours associated with high intratumoural solid stress (*12*, *38*) (Fig. 3H). A similar negative correlation between eEF2K expression and overall survival was also observed in mechanically challenged basal breast and colon cancers (Supplementary Figure 3 E,F), suggesting that eEF2 kinase activity may contribute to tumour adaptation to mechanically challenging environments. To further test this, we generated an eEF2K knockout (KO) line by CRISPR-Cas9-mediated gene editing of the transplantable murine pancreatic ductal adenocarcinoma cell line, KPC (Supplementary Fig. 3G). Several guides were tested, the sg1 clone, which showed the greatest reduction in eEF2K levels, was selected for subsequent use. Both KO and control (Rosa-targeted) cells were orthotopically injected, and tumours were harvested one-month post-injection. We then analysed eEF2 phosphorylation in the epithelial compartment of control (Rosa-targeted) KPC tumours and found that phosphorylation was most pronounced in regions positive for picrosirius red staining, suggesting that eEF2K is activated in more mechanically challenged, collagen-rich microenvironments (Supplementary Fig. 3H). In line with this, tumour measurements revealed a trend toward volume reduction in the eEF2K KO cohort, consistent with a role for eEF2K in sustaining tumour growth within the mechanically challenging PDAC microenvironment (Supplementary Fig. 3I). Together, these results identify eEF2K as a mechanosensitive effector that couples compressive stress to translational repression and tumour adaptation in PDAC.

### Early translation repression is mediated by Piezo1-dependent ER Ca²⁺ signalling

Eukaryotic elongation factor 2 kinase (eEF2K) is a member of the atypical α-kinase family and the only known calcium/calmodulin-dependent kinase within this group (*39*). eEF2K is unusual among Ca²⁺/calmodulin-dependent kinases in that binding of Ca²⁺-loaded calmodulin to its calmodulin-binding domain acts as an allosteric activator. This interaction induces a conformational change that promotes autophosphorylation of eEF2K at the activating residue T348, thereby stimulating kinase activity (*40*, *41*). Its activity is further modulated by inhibitory phosphorylation of several serines by mTORC1. Mechanical stimuli have been shown to induce Ca²⁺ flux by activating mechanosensitive ion channels, such as Piezo1, located on both the plasma membrane and the endoplasmic reticulum (*42*, *43*). Therefore, we hypothesized that compressive stress increased calcium flux to activate eEF2K. Intracellular calcium levels were monitored using the calcium indicator Fluo-8 AM. Cells were loaded with 5 µM Fluo-8 AM, baseline fluorescence was recorded, and cells were then subjected to 10 kPa compressive stress. Calcium dynamics were imaged continuously for 30 minutes following compression. We observed a significant increase in fluorescence intensity following compression, confirming Ca^2+^ release in response to application of the compressive stress used in our system (Fig. 4A).

**Figure 4.**
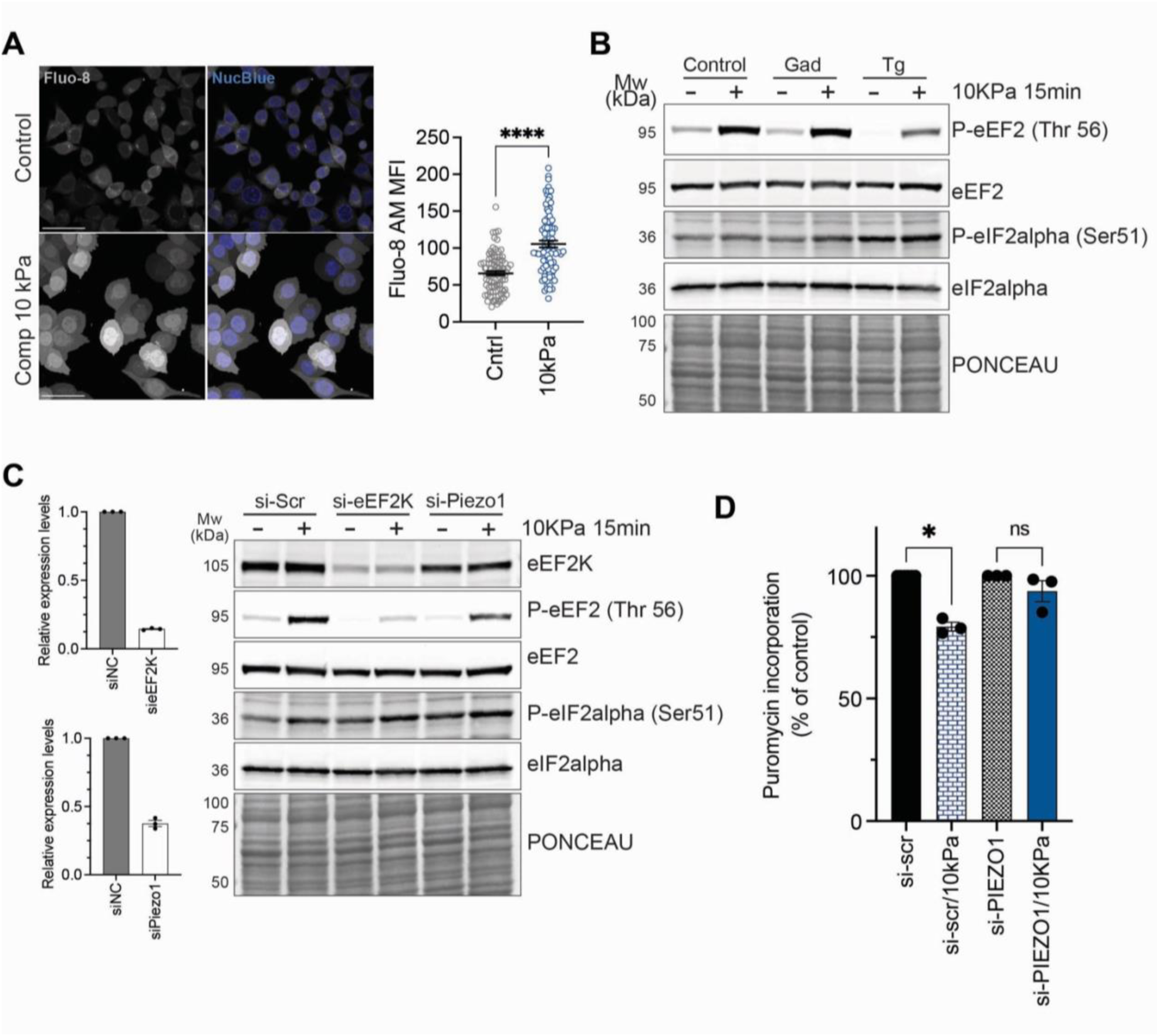
Compression induced release of ER calcium stores activate eEF2kinase to tune translation. **A.** Representative confocal image from live cell imaging of Fluo-8 loaded control and 10 kPa compressed cells. Scale bar represents 50 *μ*m. (left) Relative quantification of MFI for control(n=88) and 10 kPa compressed(n=84) cells. Data points represent individual cells plotted as Mean ± SEM, for data pooled from n=2 individual experiments. (right) **B.** Representative western blot analysis of eEF2 (T56) and eIF2α (S51) phosphorylation status in cells under control and 10 kPa compressed conditions following treatment with 25*μ*M Gadolinium or 2*μ*M Thapsigargin. Ponceau S staining serves as a loading control. Blots are representative of n=3 independent experiments. **C.** qRT-PCR of eEF2Kinase and Piezo1 mRNA levels following 24-hour siRNA treatment. Data is represented as mean ±SEM, individual points represent n=3 individual experiments(left). Representative western blot analysis of eEF2 (T56) and eIF2α (S51) phosphorylation status upon application of 10 kPa compression in control, sieEF2K and siPiezo1 cell lines. Ponceau S staining serves as a loading control. Blots are representative of n=3 independent experiments (right). **D.** Quantification of puromycin incorporation in siScrambled and siPiezo1 cells under control and 10 kPa compressed conditions. Data points represent independent experiments plotted as Mean ± SEM for n=3 independent experiments. Un-paired t-test with Welsh’s correction (A) One-way ANOVA with Sidak’s multiple comparison test (D). *P* values: ∗ *P* < 0.05, ∗∗∗∗ *P* < 0.0001.

We next investigated the source of the calcium signal using gadolinium chloride (GdCl₃) and thapsigargin (Tg). GdCl₃, a membrane-impermeant trivalent cation that inhibits a range of mechanosensitive and stretch-activated Ca²⁺ channels (*44*), did not suppress compression-induced eEF2 phosphorylation (Fig. 4B). We therefore asked whether intracellular Ca²⁺ stores contribute to the response. Thapsigargin is a specific, irreversible inhibitor of the Sarco/Endoplasmic Reticulum Ca²⁺-ATPase (SERCA) and has been used to deplete ER Ca²⁺ stores prior to mechanical stimulation (*45*)(*46*). Cells were treated with 2 µM Tg for 5 minutes, followed by washout and a 30-minute equilibration period before compression. This approach was used to deplete the releasable ER Ca²⁺ pool while minimizing continued exposure to Tg during the mechanical stimulus. Under these conditions, Tg pretreatment markedly reduced the compression-induced increase in eEF2 phosphorylation (Fig. 4B). Consistent with this, Fluo-8 imaging confirmed that Tg pretreatment significantly attenuated the compression-induced increase in intracellular Ca²⁺ (Supplementary Fig. 4A). Together, these findings indicate that an intact ER Ca²⁺ store is required for the acute activation of eEF2K in response to compression.

We then investigated the contribution of mechanosensitive Piezo channels to eEF2 phosphorylation. Piezo1 has been reported both at the level of the plasma membrane and the ER and is an established mediator of mechanically induced Ca²⁺ signalling(*47*, *48*). We first verified the subcellular localization of Piezo1 and confirmed its presence at the level of the ER and plasma membrane (Supplementary Fig. 4B). As expected, Piezo1 depleted cells did not show a significant increase in intracellular Ca^2+^ as measured by Fluo-8, under compression (Supplementary Fig. 4A). Consistent with these findings, Piezo1 depletion partially reduced compression-induced eEF2 phosphorylation (Fig. 4C). Finally, we asked whether Piezo1 knock-down would rescue the mechano-induced translational repression. In comparison to the siScramble controls, Piezo1 knockdown cells displayed a marked rescue of the compression induced translational repression (Fig. 4D). Collectively, these findings identify a Piezo1 dependent ER Ca²⁺ signaling axis upstream of eEF2K activation.

### Piezo1-linked mitochondrial dysfunction and ROS production reinforce compressive stress–induced translational repression

While depletion of Piezo1 partially rescued translation under compression, these interventions did not fully restore biosynthetic activity to baseline levels. This suggested that additional stress-responsive pathways act downstream or in parallel to the Piezo1–Ca²⁺–eEF2K axis to reinforce translational repression. Because eEF2 kinase activity is known to be regulated by oxidative stress (*49*, *50*), we hypothesized that compressive stress may induce mitochondrial dysfunction and reactive oxygen species (ROS) production that further sustain translational inhibition.

To explore this possibility, we first examined the structural and functional state of mitochondria under compression. Live-cell imaging of HCT116 cells expressing mitoDsRed revealed rapid and widespread remodelling of the mitochondrial network upon 10 kPa compression, characterized by pronounced fragmentation and the formation of perinuclear toroidal mitochondria (Fig. 5A). These morphological changes were accompanied by the assembly of filamentous actin around mitochondria, consistent with mechanically driven mitochondrial remodelling (Supplementary Fig. 5A)(*51*). Transmission electron microscopy was carried out on control and compressed samples to gain more detailed insight into the impact of compression on mitochondrial structures. The micrographs revealed striking alterations into the organization of the cristae in compressed cells (Fig. 5B, and Supplementary Video A and B), indicative of impaired inner membrane organization and compromised bioenergetic capacity.

**Figure 5.**
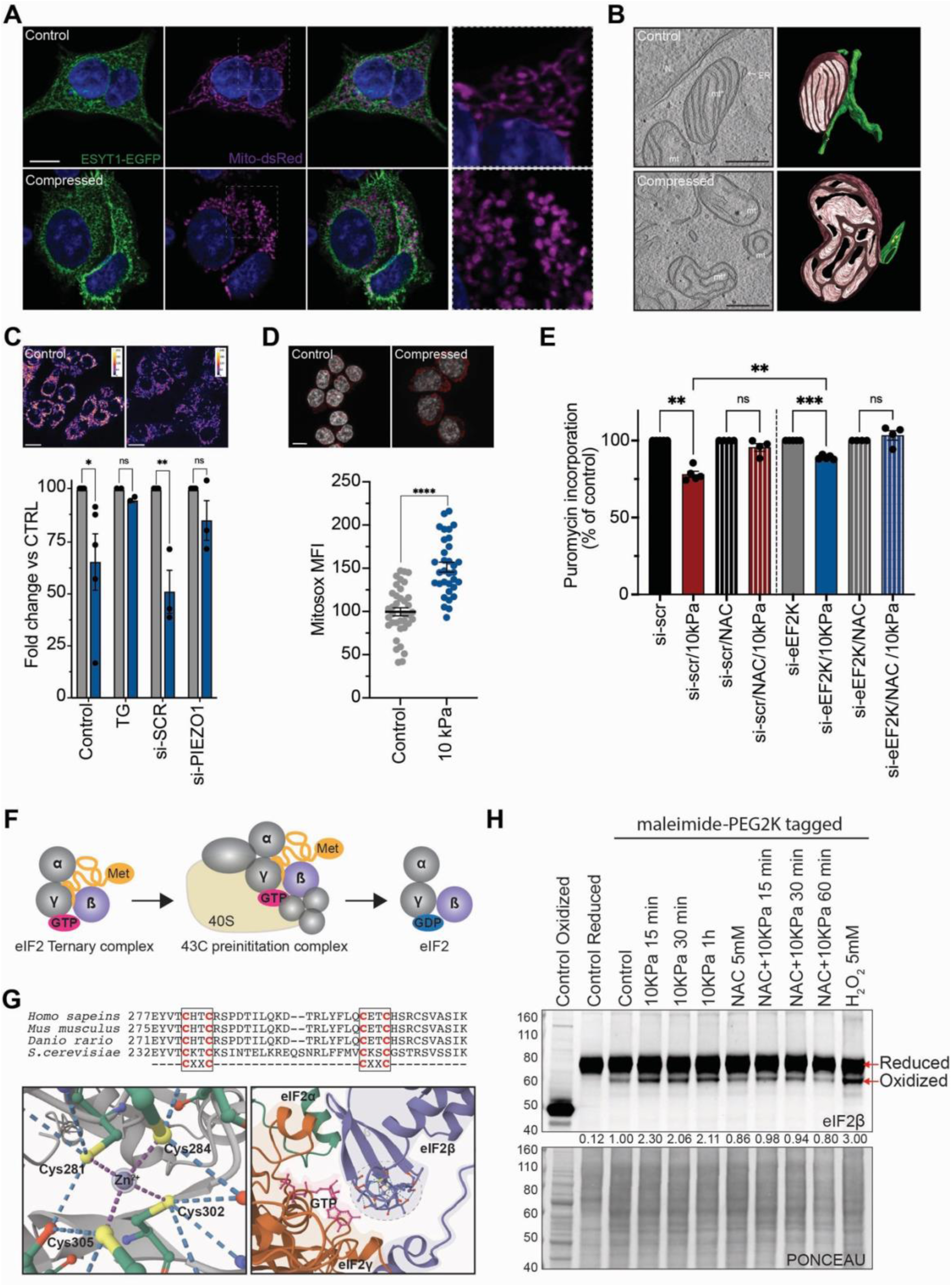
Piezo1 derived Ca2+ release drives mitochondrial ROS production to support translational repression. **A.** Representative images of mitochondrial fragmentation upon compression, with enlargement of the mitochondrial panel to highlig ht the morphological differences. ESYT-EGFP labels ER with Mito-dsred used to label mitochondria. Scale bar represents 10 *μ*m. **B.** Single tomographic slices from EM tomography reconstructions and 3D models of representative mitochondria from control and 10kPa compressed cells (60 min). Scale bar represents 400*n*m. **C.** Representative images of tetramethylrhodamine methyl ester (TMRM) incorporation in control and compressed conditions. The scale bars represents 10 *μ*m (above). FACS based quantification of TMRM incorporation normalized to total Mito-tracker Green for Control and 10kPa compressed cells for: control (n=≥3), Tg (n=2), siScr (n=3, and siPiezo1 (n=3) treated cells. Data represent independent experiments plotted as Mean ± SEM **D.** Representative imaging of Mitosox superoxide indicator for control and 10 kPa compressed cells. Scale bar represents 10 *μ*m (above). MFI for Mitosox in control (n=37) and 10kPa compressed(n=33) conditions, pooled from n=2 independent experiments. Data points represent individual cells plotted as Mean ± SEM. **E**. Puromycin incorporation in control in 10 kPa compressed ± N-acetylcysteine treatment in control and si-eEF2Kinase cells. Data is represented as Mean ± SEM for n=≥3 independent experiments. **F.** Schematic representation of role of the eIF2 complex in the formation of the 43C preinitiation complex. **G.** Sequence alignment of conserved CX_2_C motifs of eIF2*β*(top) Alphafold generated images of the Zinc coordinated tetrahedral cysteine structure of eIF2*β*(bottom left), and placement of the motif adjacent to the GTP binding pocket of the eIF2 complex. **H.** Representative western of Malemide-PEG2K tagged eIF2*β* upon compression in control, 10 kPa compressed, NAC (5mM), and H_2_O_2_ treated conditions. Ponceau S staining serves as a loading control. Blots are representative of n=3 independent experiments. One-way ANOVA with Sidak’s multiple comparison test(C,E), Unpaired t-test, two-tailed (D). *P* values: ∗ *P* < 0.05, ∗∗ *P* < 0.01, ∗∗∗ *P* <

We next assessed mitochondrial function by measuring membrane potential using tetramethylrhodamine methyl ester (TMRM), a cationic dye that accumulates in mitochondria with active membrane potential. Compressed cells displayed a clear reduction in TMRM fluorescence relative to Mitotracker Green, indicative of a loss of polarization (Fig. 5C). Mitochondria are key regulators of cellular energetics and are highly sensitive to calcium fluctuations, as excessive calcium uptake can trigger mitochondrial depolarization through opening of the permeability transition pore and disruption of the proton gradient(*52*). Given that Piezo1 activation leads to ER calcium release upon compressive stress, we hypothesized that this calcium flux may contribute to mitochondrial dysfunction. Importantly, both Piezo1 depletion and thapsigargin pretreatment, each of which interferes with ER Ca²⁺ release, significantly mitigated mitochondrial depolarization (Fig. 5C). These results identify ER-derived Ca²⁺ flux downstream of Piezo1 as a major contributor to mitochondrial dysfunction under compressive stress.

Mitochondrial impairment is tightly coupled to ROS generation. Consistent with this, compressed cells exhibited increased fluorescence of both DCFDA, a general oxidative indicator, and MitoSOX, a probe specific for mitochondrial superoxide (Supplementary Fig. 5B and Fig. 5D). Together, these findings indicate that Piezo1-mediated Ca²⁺ release disrupts mitochondrial function and elevates oxidative stress under mechanical confinement. Previous studies suggest that ROS can activate eEF2 kinase as part of a stress-adaptive translational response (*50*). To test whether this mechanism operates in our system, we examined the role of ROS in eEF2 kinase activation under compressive stress. We used *N*-acetyl-L-cysteine (NAC), a well-established ROS scavenger, to reduce intracellular ROS levels. Notably, NAC pre-treatment partially suppressed eEF2 kinase activation in response to compression (Supplementary Fig. 5C). These findings collectively suggest that Piezo1-mediated ER calcium release under compressive stress triggers mitochondrial dysfunction and ROS production, which in turn reinforce translational repression through eEF2 kinase activation. If compression-induced ROS partially regulates eEF2 kinase activity, then pretreatment with NAC would be expected to rescue translation to levels comparable to those in siEEF2K-treated cells. Surprisingly, we observed an additional rescue over that conferred by siEEF2K, suggesting that ROS may be regulating translation under compression in a complementary fashion (Fig. 5E).

Cytosolic protein biosynthesis is highly responsive to oxidative stress, and redox regulation of translational components represents a key adaptive mechanism (*53*). Among these, eIF2β, a core subunit of the trimeric eIF2 complex and central regulator of translational initiation, contains a highly conserved Cys–X₂–Cys–X₁₇–Cys–X₂–Cys motif predicted to form a zinc-coordinated tetrahedral structure at the interface of the GTP-binding pocket (Fig. 5F,G). This motif was identified as redox-sensitive in large-scale proteomic analysis of the *S. cerevisiae* redoxome (*54*), and eIF2β has since been confirmed as a zinc-binding protein in human cells (*55*), raising the possibility that oxidation of these coordinating cysteines could disrupt eIF2 function under oxidative stress.

To test whether compressive stress induces eIF2β oxidation, we employed a maleimide–PEG (2kDa) redox–Western blot assay (*56*). We detected a partially oxidized eIF2β species following compression, resembling that induced by acute H₂O₂ treatment and abolished by NAC pretreatment under the same conditions that rescued puromycin incorporation (Fig. 5H). To address the universality of this we extended the analysis to eIF5 and EEF2. Both have been previously reported to have redox sensitive cysteines, and eIF5 contains a predicted zinc-coordinated cysteine tetrahedral conformation in the structure, although it is distant from the GDP binding pocket. Neither protein showed significant oxidation upon compression or exposure to compression, suggesting that oxidation under the conditions induced by compression is confined to a distinct group of proteins (Supplementary Fig. 5 D,E). Together, these data suggest that Piezo1-dependent ER Ca²⁺ release destabilizes mitochondrial function, elevates ROS, and promotes redox-dependent oxidation of eIF2β, potentially contributing to sustained translational repression.

### AMPK-driven autophagic remodelling restores cytoplasmic density under compressive stress

Having established the acute Piezo1-Ca²⁺-eEF2K signalling axis that represses translation under compression, we next investigated how cells recover from this mechanically induced increase in macromolecular crowding reported in Figure 1C. The rapid resolution of macromolecular crowding observed in compressed cells (Fig. 1C) suggested an active recovery process, such as autophagy. We have shown that compression induces both elevated calcium flux and mitochondrial dysfunction, two convergent signals known stimulate AMPK activation (*57–60*). We therefore hypothesized that analogous to eEF2K, AMPK might be activated downstream of Ca²⁺/calmodulin-dependent protein kinase kinase β (CAMKKβ) to trigger autophagic remodelling and restore cytoplasmic density homeostasis independently of mTORC1. To test this, we analyzed the kinetics of autophagy onset following compression by monitoring AMPK Thr172 phosphorylation, AMPK substrate Unc 51-like kinase 1 (ULK1) Ser555 phosphorylation and LC3B lipidation. Phosphorylation of AMPK at Thr172 exhibited kinetics comparable to those of eEF2K (Fig. 6A). This was accompanied by phosphorylation ULK1 at Ser555 and LC3B lipidation at 30 minutes (Fig. 6A). Notably, AMPK has recently been recognized as a mechano-responsive kinase, activated in response to mechanical stimuli such as those occurring during interstitial cell migration, a context in which cells experience significant compressive stress (*61*).

**Figure 6.**
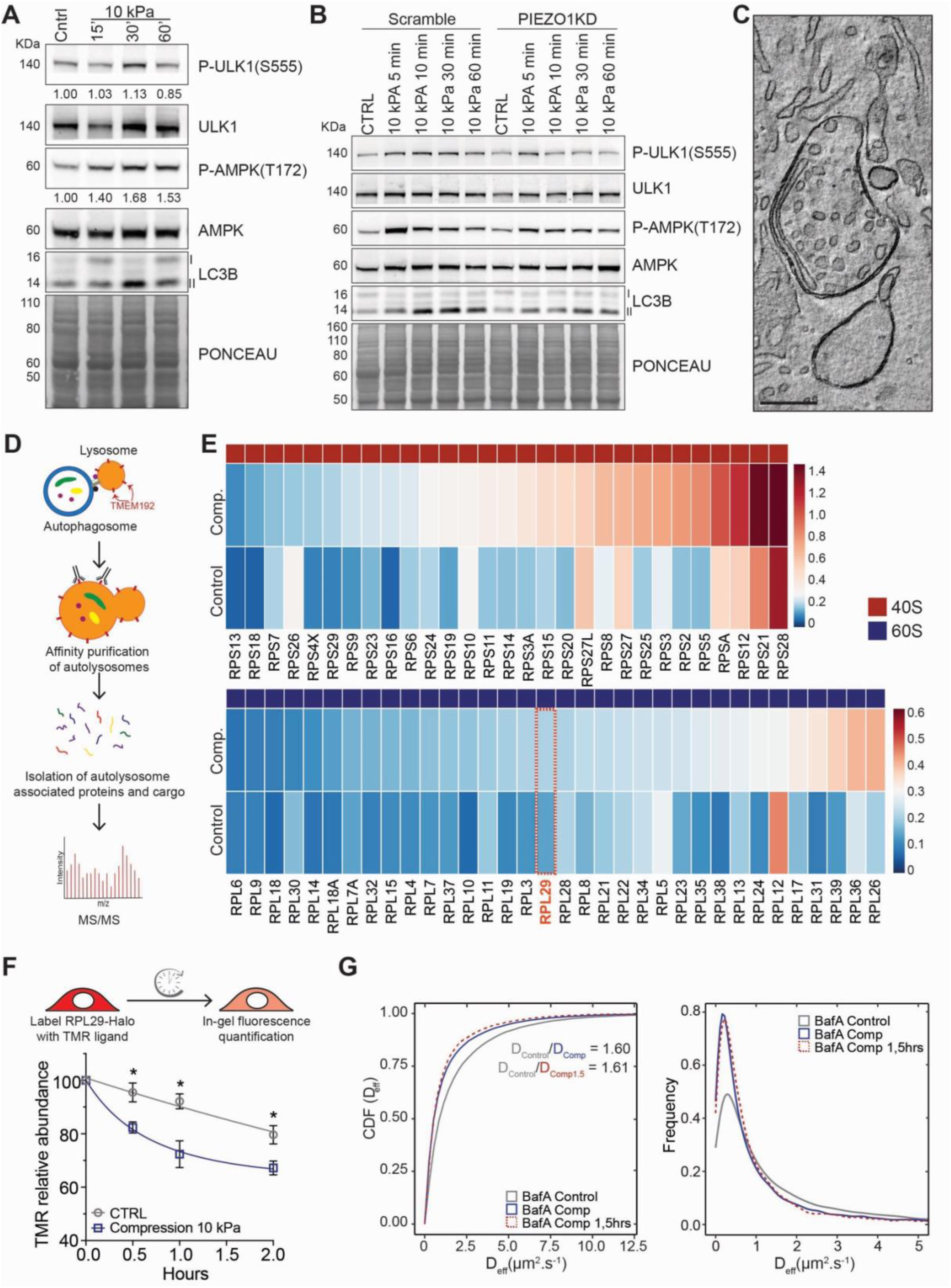
Ribostasis regulates rheological homeostasis under acute mechanical stress. **A.** Western blot analysis of kinetics of AMPK activation, ULK1 Ser555 phosphorylation and LC3B lipidation(II) following 10kPa compression. Ponceau S staining serves as a loading control. Representative of n=2 independent experiments **B**. Western blot analysis of compression activation of AMPK and autophagy markers: ULK1 phosphorylation and LC3B lipidation in control and Piezo kd cells. Ponceau S staining serves as a loading control. Representative of n=2 independent experiments **C.** A single tomographic slice from EM tomography reconstruction of autophagosomes in 30-minute compressed sample. Scale bar represents 200 *n*m. **D.** Schematic of Tmem192-3HA pulldown used to enrich for lysosome and autophagosome populations **E.** Percent enrichment of ribosomal proteins in TMEM192-immunopurified lysosomal fractions from control and 10 kPa compressed (1 h) HCT116 cells, normalized to whole-cell extract and quantified by mass spectrometry. Heatmaps show individual ribosomal proteins ranked by relative enrichment under compression; 40S (red) and 60S (blue) subunit proteins are indicated. **F.** Quantification of in-gel fluorescence of RPL29-HALO degradation kinetics following an initial labeling with HALO ligand TMR. Data is represented as Mean ±SEM for n=3 independent experiments. **G.** Cumulative distribution function showing effective diffusion co-efficient (D_eff_) data for 40nM GEMs in Bafilomycin A (200nM) treated HCT116 for control (grey), compressed 0-30 minute (blue), and compressed 1.5 hr (red) conditions and distribution of 40nM GEM D_eff_ measured in control (grey), compressed 0-30 minute (blue), and compressed 1.5 hr (red) conditions. Data represents pooled data from 2 independent experiments. 2-way ANOVA with Sidak’s multiple comparison test(F). *P* value: ∗p < 0.05

To assess the calcium dependency and the role of Piezo1 on the activation of autophagy we examined AMPK Thr172 phosphorylation, ULK1 Ser555 phosphorylation, and LC3 lipidation in control and Piezo1kd cells. In control cells we observed a rapid and transient increase in AMPK Thr172 and ULK1 Ser555 phosphorylation accompanied by enhanced LC3 lipidation. In contrast these responses were abolished in the Piezo1kd cells (Fig. 6B). These data suggested that similar to EEF2K activation, AMPK activation occurs downstream of compression induced Piezo1-mediated Ca^2+^ influx, which drives activation of calcium responsive kinases to regulate proteostasis. This response was further supported by a significant increase in, both the number and size of p62 foci (Supplementary Fig. 6A). Consistent with the biochemical findings, transmission electron microscopy revealed well-defined autophagosomes in cells subjected to 10 kPa compression for 30 minutes (Fig. 6C). The rapid LC3B lipidation we observed preceded the observed delayed inactivation of mTORC1 signaling under compression (Fig. 2B). To further demonstrate the mTORC1 independence of the autophagic response, we examined LCB-II emergence in TSC2KD cells. LC3B-II accumulated with similar kinetics in both TSC2KD and control cells, confirming that compression-induced autophagy proceeds independently of mTORC1 (Supplementary Fig. 6B). Collectively, these findings indicate that Piezo1-mediated Ca^2+^ influx and subsequent AMPK activation trigger an autophagic response following compression, prompting us to investigate whether this process is coupled to enhanced protein turnover and selective regulation of specific protein compartments.

To assess whether compression leads to increased protein loss, we labeled newly synthesized proteins with L-azidohomoalanine (AHA), a clickable methionine analogue that is incorporated into newly synthetized proteins without cessation of translation. Following labeling with AHA we monitored AHA-containing protein levels over a three-hour period under control and 10 kPa compressed conditions. We observed a significant increase in overall protein loss in compressed versus control conditions (Supplementary Fig. 6C). To determine whether specific classes of proteins were being targeted, we engineered an HCT116 cell line expressing Tmem192-3HA to enable affinity-based lysosome capture followed by mass spectrometry identification of lysosomal and autophagosomal cargo (Fig. 6D) (*62*, *63*). Lysosome-enriched proteins were normalized to whole cell protein to identify proteins that were selectively accumulated in lysosomal fractions. Comparison of GO enrichment profiles in compressed and control cells revealed an over-representation of translation-related proteins (Supplementary Dataset 2 and Supplementary Fig. 6D). Notably, ribosomal proteins were strongly enriched in the Tmem192 fractions of compressed cells (Fig. 6E). The enrichment was more pronounced for ribosomal proteins pertaining to the 40S subunit, though several 60S proteins, including RPL29, were also enriched in the lysosomal fraction under compression (Fig. 6E). To validate these findings, we employed the HCT116 Ribo-HALO cell line directly measure degradation rates of RPL29 under compression. Cells were pulse-labeled with HALO ligand TMR for one hour, washed to remove unbound ligand, and then subjected to 10 kPa compression or maintained under control conditions. In-gel fluorescence analysis of labelled RPL29 revealed a rapid and significant loss of RPL29 beginning at 30 minutes post-compression, corroborating the mass spectrometry results and supporting the conclusion that compression induces a ribosomal degradation (Fig. 6F). Previous work has shown that NUFIP1 (nuclear fragile X mental retardation-interacting protein 1) translocation from the nucleus to the cytosol directs ribosomal proteins for degradation under nutrient deprivation(*63*). We therefore examined whether mechanical stress was able to induce a similar redistribution of NUFIP1 to the cytosol to direct ribophagy programs. However, NUFIP1 was not present in lysosomal fractions (Supplementary Dataset 2), nor did we observe a significant cytosolic enrichment of NUFIP1 by immunofluorescence (Supplementary Fig. 6E), suggesting that the ribophagy program induced following compression proceeds through an alternative mechanism. Given the tight coupling of ribosome biogenesis to the translational activity of the cell, we next asked whether ribosome production was disrupted in compressed cells by quantifying 47S rRNA levels(*64*). Quantitative real-time PCR using primers targeting the 5’ region of the A’ splice site revealed a marked reduction in 47S rRNA levels beginning at 60 minutes after compression onset (Supplementary Fig. 6F). These data indicate that compressive stress specifically disrupts ribosome homeostasis, impacting both degradation and biogenesis pathways.

Finally, ribosomes have been proposed to be one of the major contributors to macromolecular crowding, as measured by the 40nM GEMs (*25*). Therefore, we asked whether the compression induced autophagy program contributed to the observed recovery of the rheological properties under sustained stress. GEM diffusion measurements under compression in the presence of autophagy inhibitor Bafilomycin A. Cells were pre-treated with 200 nM Bafilomycin A for two hours, prior to initiating compression. 10kPa of compression were applied to the cells and measurements were made at 0-30 minutes and 90-120 minutes. Bafilomycin A treated cells exhibited both a more pronounced increase in baseline macromolecular crowding compared to control cells and a markedly impaired ability to reestablish baseline values under continued compression (D_cntrl_=0.874 µm^2^s^-1^, D_10kPa/0min_=0.543 µm^2^s^-1^, D_10kPa/90min_=0.542 µm^2^s^-1^)(Fig. 6G). These findings suggest that the reestablishment of cytoplasmic density homeostasis under mechanical stress is achieved in part by active remodeling of ribosome homeostasis.

## Discussion

Cells exist within mechanically dynamic environments, yet how they preserve intracellular organization and proteostatic balance under physical compression has remained unclear. Our findings reveal that compression does not act as a single uniform break on protein synthesis but instead engages three mechanistically distinct Ca^2+^ dependent arms: an eEF2K dependent tuning of elongation, AMPK-driven ribophagy and ROS-dependent oxidation of eIFβ, that together adapt proteostatic homeostasis to the mechanically challenged environment (Fig. 7). This architecture is not redundant so much as combinatorial, each arm targets a different step (translation initiation, elongation, and ribosome abundance) and operates on different timescales, allowing the cell to tune biosynthetic output proportionally to the severity and duration of the mechanical load rather than executing and all or nothing shutdown. Consistent with this model, disrupting Piezo1, produces a translational rescue (Fig. 4D) disproportionally larger than its effect on eEF2K activation alone (Fig. 4C), because Piezo1 sits upstream of both the eEF2K and ROS/eIF2β arms (Fig. 5E). We propose that this multi-armed, Piezo1-nucleated design allows cells to adapt their translational response to a compressive load in a way that a single linear pathway could not.

**Figure 7.**
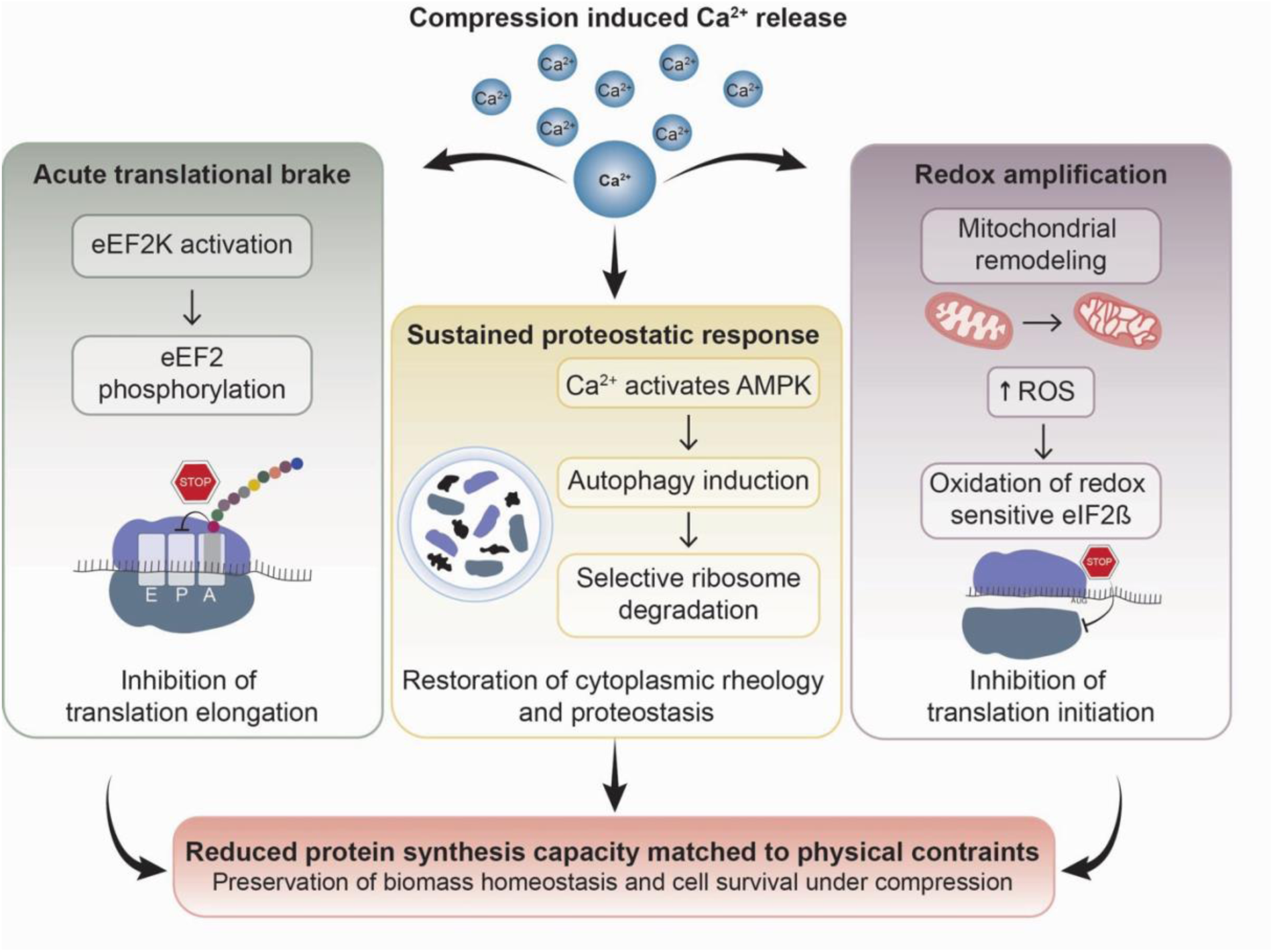
Compression-induced Ca²⁺ release coordinates a multi-armed translational stress response to preserve biomass homeostasis. Mechanical compression triggers Ca²⁺ release, which engages three parallel effector arms. Left: Ca²⁺ activates eEF2K, leading to eEF2 phosphorylation and acute slowing of translation elongation. Center: Ca²⁺-dependent AMPK activation induces autophagy and selective ribosome degradation (ribophagy), restoring cytoplasmic rheology and proteostasis. Right: compression-associated mitochondrial remodelling elevates ROS, which oxidizes redox-sensitive eIF2β to inhibit translation initiation. Together, these mechanisms reduce protein synthesis capacity in proportion to the physical constraints imposed by compression, preserving biomass homeostasis and cell survival.

Our findings further indicate that the early adaptive response to compression proceeds largely independently of mTORC1 and the integrated stress response, despite their canonical roles in stress-responsive translational control(*18*, *21*, *25*). This independence is not simply a matter of degree: compression-induced macromolecular crowding, even though measurably lower than that produced by mild osmotic stress (0.1 M sorbitol), left mTORC1 signalling acutely intact over a timescale in which osmotic stress rapidly and completely inactivated it. A purely diffusion-limited model of translational control, in which crowding above a critical threshold restricts ribonucleoprotein or vesicle mobility regardless of its origin (*65*, *66*), would predict the opposite. We therefore favour a model in which osmotic and compressive stress are sensed through distinct routes: osmotic shock likely restricts vesicular trafficking at a scale sufficient to destabilize mTORC1 signalling directly, whereas the compression regimes used here, comparable to physiological solid stress in tumours (*12*, *13*), remain below this threshold and instead engage ER-localized Piezo1 to trigger Ca²⁺-dependent signalling without immediately perturbing mTORC1. Notably, mTORC1 activity was not permanently insulated from compression: even TSC2-depleted cells showed a delayed loss of mTORC1 target phosphorylation at later time points, indicating a TSC2-independent route to eventual mTORC1 silencing whose identity remains an open question for future work.

A second point warranting discussion is the redox regulation of eIF2β itself. Zinc-coordinated cysteine motifs of the kind we identify in eIF2β have been described as redox switches in a handful of other translation factors (*53–56*), but functional evidence linking their oxidation to a physiological challenge, as opposed to exogenous oxidant exposure, has been limited. Our finding that compression induces eIF2β oxidation through a Piezo1–mitochondrial ROS axis, at a magnitude resembling that induced by direct H₂O₂ treatment and reversible by NAC, extends this class of redox-regulated translation factors to a physiologically relevant mechanical trigger. Because eIF2β oxidation acts at the level of translation initiation while eEF2K acts at elongation, this provides a biochemical basis for why loss of either mechanism alone only partially rescues translation: the two operate on independent steps of the translation cycle and are only redundant in their downstream effect, not in their molecular target.

Beyond acute repression, we find that cells actively resolve compression-induced crowding that arises through volume reduction through AMPK-dependent, selective ribosome degradation, restoring GEM diffusivity to baseline within 90 minutes despite sustained mechanical load. This recovery required autophagic flux, as Bafilomycin A treatment both exaggerated baseline crowding and prevented its resolution. While recent work has emphasized polysome assembly and dissolution as direct modulators of cytoplasmic rheology (*19*, *20*), our results suggest a complementary, slower-acting mechanism in which the cell actively remodels its ribosome population to relieve compression-induced crowding. Notably, this response is accompanied by a reduction in 47S rRNA levels beginning at approximately 60 minutes after compression, indicating that sustained mechanical stress affects not only ribosome turnover but also ribosome production. Together, these findings suggest that compression remodels ribosome homeostasis at multiple levels, coupling selective ribosome clearance with reduced rRNA abundance as translational demand remains suppressed. The delayed reduction in 47S rRNA relative to the rapid activation of eEF2K is consistent with a temporal hierarchy in which acute mechanical stress first limits translation, followed by broader adjustment of ribosome abundance during sustained compression. Notably, the ribophagy response proceeds independently of NUFIP1 re-localization (*63*), pointing to an alternative, as-yet-undefined route into ribosome clearance under mechanical rather than nutrient stress.

These findings may be relevant to how solid tumours cope with mechanical contraing. Intratumoural solid stress is an established driver of therapy resistance and metabolic reprogramming (*10–14*). Surveying publically available datasets we have shown that elevated eEF2K expression associates with poor clinical outcomes specifically in pancreatic ductal adenocarcinoma, one of the most mechanically stressed solid tumours (*12*, *67*), with a similar trend evident in basal breast cancer, and to a lesser degree in colon cancer. Together with our observation of a trend towards reduced tumour growth following orthotopic injection of KPC cell lines with eEF2K KO, these findings are consistent with a potential role for eEF2K in tumour adaptation to mechanical constraint.

Several aspects of our study provide clear directions for future work. Our compression system, while tunable and physiologically calibrated, applies stress over a defined 2D geometry and timescale that cannot fully recapitulate the heterogeneous, sustained stress fields of a growing tumour mass. The eEF2K-knockout orthotopic tumour data reported here are limited in cohort size and should be regarded as a preliminary indication of functional relevance requiring further validation rather than definitive evidence of a growth-restricting phenotype. Finally, the TSC2-independent, delayed mTORC1 inactivation we observe under sustained compression remains mechanistically undefined and represents a natural extension of this work. Addressing these points, together with defining the precise biophysical parameters that distinguish mechanically-from osmotically-induced crowding, will be important next steps in establishing the generality of this checkpoint across mechanically diverse tumour contexts.

## Acknowledgements

This research was supported by Project: 101142614 — TOPOMECH — ERC-2023-ADG. Giuseppe Fragale was supported by an AIRC Pre-doctoral Fellowship Ref. ID 28149 and a 2-year AIRC Post-doctoral Fellowship Ref. ID 29811. Yuwei Li was supported by an AIRC Pre-doctoral Fellowship Ref. ID 33063. MCL was supported by an AIRC (Associazione Italiana per la Ricerca sul Cancro) IG 2019 grant ID. 23258 and IG 2024 grant ID. 30391. PM was supported by an AIRC (Associazione Italiana per la Ricerca sul Cancro) IG grant ID. 24976 and the Italian Ministry of University and Scientific Research, PRIN PNRR P2022F3YRF. We thank Angela Cattaneo for technical support for the phospho-proteomics. We thank Marie Albinet, Ruchika Kumari, Marco Foiani, Simona Polo and Giorgio Scita for discussion and feedback on the manuscript.

## Author Contributions

K.M.H. conceived and designed this study. G.F. performed the majority of the experiments, conducted data analysis, and contributed to figure editting. B.J.H, Y.L, W.X., N.B., N.M., and P.M. provided experimental assistance and participated in data analysis. G.P., G.V.B, E.A., Z.L., V.M., F.P., and D.M., provided technical assistance and participated in data analysis. A.A. contributed reagents and provided experimental support. B.J.H., L.M., A.K., M.G., and A.E.M. provided technical assistance in support KPC cell line construction and in vivo studies. D.P, A.B. and M.C.L. provided technical supervision and constructive suggestions. K.M.H. wrote the manuscript with comments from all authors.

## Materials and Methods

### Cell lines

HCT-116 (DSMZ ACC 581) cells and derivative HCT-116 TSC2KD cells were cultured in RPMI 1640 medium without L-glutamine (Euroclone, Cat# ECB9006L) supplemented with 10% fetal bovine serum (FBS; Euroclone, Cat# ECS5000L) and 1% L-glutamine (Euroclone, Cat# ECB3000D). Human embryonic kidney 293 cells expressing the simian virus 40 (SV40) large T antigen (HEK293T; ICLC) were cultured in DMEM high glucose (Euroclone, Cat# ECB7501L) supplemented with 10% Fetal Bovine Serum (FBS). hTERT-RPE1 (ATCC CRL-4000) cells were cultured in DMEM/F-12, GlutaMAX™ (Life Technologies, Cat# 31331-098) supplemented with 10% FBS and 0.5% sodium pyruvate (Biowest/VWR, Cat# L0642-100). All cell lines have been authenticated by cell fingerprinting and tested for mycoplasma contamination. Cells were grown at 37 °C in a humidified atmosphere with 5% CO2.

### Plasmids

The pLKO.1-TSC2 plasmid was a gift from Do-Hyung Kim (Addgene plasmid #15478; RRID: Addgene_15478) (*68*).The pLKO.1-hygro plasmid was a gift from Bob Weinberg (Addgene plasmid #24150; RRID: Addgene_24150). The pcDNA3-TORCAR plasmid was a gift from Jin Zhang (Addgene plasmid #64927; RRID: Addgene_64927). For expression of the 40-nm genetically encoded multimeric nanoparticle (GEM), pCMV-PfV-Sapphire-IRES-DSRed was a gift from Liam Holt (Addgene plasmid # 116934; http://n2t.net/addgene:116934; RRID:Addgene 116934) was used as described in (*25*). The pX459 plasmid used to assemble the RPL29-HaloTag CRISPR knock-in vector was a gift from J. Wade Harper and was used as described previously(*28*). pLJM1-Tmem192-mRFP-3xHA was a gift from Roberto Zoncu (Addgene plasmid # 134631; http://n2t.net/addgene:134631; RRID: Addgene_134631)(*69*).

### Plasmid transfection

HCT116 cells were transiently co-transfected with PIEZO1-Halo and ESYT-1-EGFP plasmids using Lipofectamine 2000 (Thermo Fisher Scientific, Cat# 11668019) in Opti-MEM I GlutaMAX™ (Life Technologies, Cat# 51985-042), following the manufacturer’s instructions and as previously described. For each well of a 6-well plate, 1.6 µg ESYT-1-EGFP and 2.4 µg PIEZO1-Halo plasmid DNA were diluted together in 250 µL Opti-MEM. In parallel, 6 µL Lipofectamine 2000 was diluted in 250 µL Opti-MEM. The two solutions were combined and incubated for approximately 15 min at room temperature before being added to the cells. The transfection medium was subsequently replaced with fresh complete medium lacking transfection reagent, and cells were used for the indicated experiments the following day.

### Antibodies

Primary and secondary antibodies used in this study are listed in the Key Resources Table. Primary antibodies included antibodies against 4EBP1, phospho-4EBP1 (Thr37/46), AKT, phospho-AKT (Ser473), AMPK, phospho-AMPK (Thr172), DRP1, phospho-DRP1 (Ser637), eEF2, phospho-eEF2 (Thr56), eEF2K, EIF2α, phospho-EIF2α (Ser51), EIF2S2 (eIF2β), eIF5, G3BP1, GAPDH, IRE1α, phospho-IRE1α (Ser724), LC3B, p38, phospho-p38 (Thr180/Tyr182), p70S6K, phospho-p70S6K (Thr389), PERK, phospho-PERK (Thr980), ULK1, phospho-ULK1 (Ser457), phospho-ULK1 (Ser555), TSC2 and vinculin. For immunofluorescence experiments, antibodies against DCP1A and G3BP1 were used. Alexa Fluorophore-conjugated secondary antibodies were used for fluorescence detection, whereas horseradish peroxidase (HRP)-conjugated secondary antibodies were used for western blotting.

### Lentivirus production

Lentiviral particles were produced in HEK293T cells using calcium phosphate transfection. HEK293T cells (5 × 10^6) were seeded in 10-cm dishes in complete DMEM without antibiotics and transfected at the time of seeding. For each dish, 10 µg of the indicated lentiviral vector (pLKO.1-hygro empty vector, pLKO.1-hygro shTSC2 or pLJM1-Tmem192-mRFP-3xHA) were mixed with 6.5 µg psPAX2 (Addgene plasmid #12260), 3.5 µg pMD2.G (Addgene plasmid #12259) and 62.4 µL 2 M CaCl₂. The DNA solution was brought to 500 µL with 0.1× TE buffer and added dropwise to 500 µL 2× HBS (pH 7.05) with gentle bubbling. After 5 min at room temperature, the transfection mixture was added to the freshly seeded cells and incubated overnight. The following day, the medium was replaced with 5.5 mL fresh complete DMEM. After a further 24 h, virus-containing supernatant was collected, filtered through a 0.45-µm filter and either used immediately for infection or stored at −80 °C.

### shRNA oligonucleotide design and vector cloning

The shTSC2 construct was generated by cloning the TSC2-targeting sequence derived from pLKO.1-puro shTSC2 (Addgene plasmid #15478) into the pLKO.1-hygro backbone following the https://www.addgene.org/protocols/plko/ protocol. The TSC2-targeting sequence was 5′-CACTGGCCTTGGACGGTATTG-3′.

### Generation of stable HCT116 shTSC2 cells

HCT116 cells (5 × 10^5 cells per well) were seeded in 6-well plates in 1 mL complete RPMI without antibiotics and infected with 1 mL shTSC2 lentiviral supernatant in the presence of 8 µg/mL polybrene (hexadimethrine bromide; Sigma-Aldrich, Cat# H9268). A second round of infection was performed the following day. After a further 24 h, the viral medium was replaced with complete growth medium containing 300 µg/mL hygromycin B (Life Technologies, Cat# 10687-010). Cells were selected for 7 days, with selective medium containing fresh hygromycin renewed three times during the selection period. Cells were passaged as required to maintain sub-confluent cultures. Following selection, 0.5–1 × 10^6 cells were lysed and TSC2 downregulation was verified by western blotting.

### siRNA transfection

HCT116 cells were reverse-transfected with siRNAs targeting eEF2K or PIEZO1, or with a non-targeting control siRNA, using Lipofectamine RNAiMAX Transfection Reagent (Thermo Fisher Scientific, Cat# 13778150). For each well of a 6-well plate, siRNA and 6 µL RNAiMAX were diluted in 500 µL Opti-MEM I GlutaMAX™ (Life Technologies, Cat# 51985-042) and incubated for 15 min at room temperature to allow complex formation. The complexes were then added to 7 × 10^5 cells seeded in 2 mL complete medium, for a final volume of 2.5 mL. siRNAs were used at a final concentration of 20 nM. After 24 h, the transfection medium was replaced with fresh complete medium, and cells were allowed to recover for several hours before being subjected to the indicated experimental treatments, including compression.

### RNA isolation and RT–qPCR

Total RNA was isolated using the RNeasy Mini Kit (Qiagen, Cat# 74104) according to the manufacturer’s instructions. For cDNA synthesis, 2 µg total RNA were reverse transcribed using the High-Capacity cDNA Reverse Transcription Kit (Thermo Fisher Scientific, Cat# 2902953). cDNA was diluted 1:40, and quantitative PCR reactions were prepared in technical triplicate using LightCycler 480 SYBR Green I Master Mix (Roche, Cat# 04707516001) in a final volume of 20 µL. qPCR was performed using a Roche LightCycler 96 instrument. Relative gene expression was calculated using the ΔΔCt method with GAPDH as the reference gene.Primer sequences were as follows: 47S rRNA, forward 5′-GCCTTCTCTAGCGATCTGAGAG-3′ and reverse 5′-CCATAACGGAGGCAGAGACA-3′; GAPDH, forward 5′-CTCAAGATCATCAGCAATGCC-3′ and reverse 5′-CATCACGCCACAGTTTCCC-3′; PIEZO1, forward 5′-CAATGAGGAGGCCGACTACC-3′ and reverse 5′-GCACTCCTGCAGTTCGATGA-3′; eEF2K, forward 5′-GCACAGCCAGAGTTACTGGAC-3′ and reverse 5′-CTCTGCCAACTCCTCTGGAC-3′.

### Drug treatments

Unless otherwise indicated, pharmacological treatments were performed in complete culture medium under standard culture conditions. For compression experiments, compounds that were maintained during treatment were added both to the culture medium and to the equilibration medium of the polyacrylamide gel overlays, so that the same final concentration was present throughout the compression step. Vehicle-treated controls (DMSO or PBS, as appropriate) were included in parallel for all experiments. Gadolinium(III) chloride hexahydrate (25μM in PBS; Sigma-Aldrich, Cat# 203289) and BAPTA-AM (10 μM in DMSO; MedChemExpress, Cat# HY-100545) were added 15 min before compression and maintained throughout the experiment. N-acetyl-cysteine (NAC; 5 mM in PBS; Sigma-Aldrich, Cat# A9165) was generally added 2 h before compression and maintained during compression. Thapsigargin (2 μM in DMSO; Selleckchem, Cat# S7895) was used as a short pre-treatment: cells were incubated for 5 min, the drug was then removed, and cells were allowed to recover for 30 min before initiation of the experimental condition (control or compression). In SUnSET assays, puromycin (Vinci-Adipogen, Cat# AG-CN2-0078-M100) was added together with sorbitol (PanReac AppliChem, Cat# A4992), water or compression, as indicated in the corresponding assay section.

### Protein extraction

Unless otherwise specified, cells were washed once with 1× PBS and lysed in 1× RIPA buffer (see Key Resources Table; 100 µL per 10^6 cells). Lysates were incubated on ice for 10–15 min and sonicated at 4 °C using a Bioruptor bath sonication system (Diagenode) for 3–5 cycles of 30 s on/30 s off. Lysates were clarified by centrifugation at 14,000 rpm for 10 min at 4 °C, and the supernatant was collected. Protein concentration was determined using the Bradford assay (Bio-Rad Protein Assay Dye Reagent Concentrate, Cat# 5000006). RIPA extracts were either stored at −80 °C or mixed with 4× Laemmli sample buffer at a 3:1 lysate-to-buffer ratio to obtain a final 1× concentration for subsequent immunoblotting.

### Immunoblotting

Primary antibodies used for immunoblotting included antibodies against 4EBP1, phospho-4EBP1 (Thr37/46), AKT, phospho-AKT (Ser473), AMPK, phospho-AMPK (Thr172), DRP1, phospho-DRP1 (Ser637), eEF2, phospho-eEF2 (Thr56), eEF2K, eIF2α, phospho-eIF2α (Ser51), eIF2β, eIF5, G3BP1, GAPDH, IRE1α, phospho-IRE1α (Ser724), LC3B, p38, phospho-p38 (Thr180/Tyr182), p70S6K, phospho-p70S6K (Thr389), PERK, phospho-PERK (Thr980), ULK1, phospho-ULK1 (Ser457), phospho-ULK1 (Ser555), TSC2 and vinculin. Antibody suppliers, catalogue numbers and working dilutions are provided in the Key Resources Table. Protein samples were denatured at 75 °C for 5–10 min before electrophoresis. Proteins were resolved on precast gradient gels (Bolt Bis-Tris Plus Mini Protein Gels, 4–12%, Invitrogen) and transferred to Amersham™ Protran 0.45-µm nitrocellulose membranes (Cytiva, Cat# 10600002) using a Mini Trans-Blot Cell system (Bio-Rad, Cat# 1703930) at 100 V for 1–1.5 h in 1× transfer buffer containing 15–20% methanol, depending on protein molecular weight. Transfer efficiency and protein loading were assessed by Ponceau staining. Membranes were blocked in TBS-T containing 5% BSA for 1 h at room temperature or overnight at 4 °C. Primary antibodies were diluted in TBS-T containing 1% BSA and incubated for 1 h at room temperature or overnight at 4 °C. After three 10-min washes in TBS-T, membranes were incubated for 1 h at room temperature with HRP-conjugated anti-mouse or anti-rabbit secondary antibodies. Following three additional washes, signals were detected using enhanced chemiluminescence and imaged using either a ChemiDoc system (Bio-Rad) or an iBright instrument (Thermo Fisher Scientific).

Densitometric quantification of immunoblots was performed using Fiji/ImageJ. Regions of interest (ROIs) of identical dimensions were applied across all lanes within the same experiment to quantify individual bands or, for SUnSET assays, the puromycin-labelled smear. Ponceau-stained total protein was quantified separately in the corresponding lanes and used for normalization. For phospho/pan analyses performed on separate membranes, phospho-specific and total-protein signals were quantified independently, and phospho-specific signals were normalized to the corresponding Ponceau signal.

### Polyacrylamide gel overlays for 1D-compression

Polyacrylamide gels and 1D overlays were prepared as previously described by Le Berre et al..(*22*, *23*) Briefly, polyacrylamide gels were polymerized on 28-mm-diameter glass coverslips. Coverslips were first cleaned with 70% ethanol and functionalized by plasma cleaning for 1 min (Diener Femto), followed by incubation in 2% (3-aminopropyl)triethoxysilane (APTES) in 95% ethanol for 15 min. Coverslips were then rinsed in 95% ethanol and dried at room temperature. Polyacrylamide gel solutions were prepared as 1-mL mixtures in 1× PBS, as reported in the table below:

| Young's modulus (KPa) | Acrylamide 40% (μL) | Bis-acrylamide 2% (μL) | APS 10% in H<sub>2</sub>O (μL) | TEMED (sigma) pure (μL) | PBS 1X (up to 1ml) (μL) |
| --- | --- | --- | --- | --- | --- |
| 2 | 125.0 (5%) | 35.0 (0.07%) | 10 | 1 | 829 |
| 5 | 185.5 (7.5%) | 30.0 (0.06%) | 10 | 1 | 773.5 |
| 10 | 185.5 (7.5%) | 47.5 (0.95%) | 10 | 1 | 756.5 |
| 15 | 185.5 (7.5%) | 65.0 (0.13%) | 10 | 1 | 738.5 |
| 20 | 185.5 (7.5%) | 100.0 (0.2%) | 10 | 1 | 703.5 |
| 30 | 250.0 (10%) | 100.0 (0.2%) | 10 | 1 | 639 |

Immediately after addition of TEMED, 200 μL of gel solution was pipetted onto a salinized glass plate, and a functionalized coverslip was placed on top. After polymerization (approximately 10 min), coverslips were detached from the parafilm with a scalpel and hydrated in PBS for 30 min. Gels were then rehydrated in PBS for approximately one hour and then equilibrated overnight at 4 °C in cell culture medium, supplemented with the appropriate drug when required, before use.

### SUnSET assay

SUnSET assays were performed as described previously (*70*). Briefly, cells were treated with 3 µg/mL puromycin (Vinci-Adipogen, Cat# AG-CN2-0078-M100) in the presence or absence of osmotic shocks or compressive stress. Untreated cells and cells treated with cycloheximide (CHX, 10 µM) during the indicated treatment period were used as negative controls. After 30 min of puromycin incorporation, cells were collected and processed for western blot analysis using an anti-puromycin antibody. Whole-lane intensity was quantified in Fiji/ImageJ and normalized to total protein determined by Ponceau staining.

### Polysome profiling

Cells were maintained in complete medium or subjected to hyperosmotic stress (0.14 M sorbitol), hypo-osmotic stress (25% H₂O), or 10 kPa compressive stress for <5 min. Immediately after treatment, cells were incubated with cycloheximide (100 µg/mL) for 10 min to arrest translating ribosomes on mRNAs. Cells were then washed with ice-cold PBS containing cycloheximide (100 µg/mL) and lysed in polysome lysis buffer containing 50 mM Tris-HCl (pH 7.5), 100 mM NaCl, 30 mM MgCl₂, 0.1% NP-40, protease inhibitors, 40 U/mL RNasin (Promega) and 100 µg/mL cycloheximide. Lysates were clarified by centrifugation at 12,000 × g for 10 min at 4 °C. Cytoplasmic extracts corresponding to 2–10 absorbance units were loaded onto 15–50% linear sucrose gradients and ultracentrifuged at 4 °C in an SW41 Ti swinging-bucket rotor (Beckman Coulter) at 40,000 rpm for 3 h 30 min. Polysome profiles were obtained by continuous monitoring of absorbance at 254 nm using a BioLogic LP chromatography system (Bio-Rad) at a flow rate of 1 mL/min(*71*).

### Nuclear volume measurements

HCT116 cells were seeded at 4 × 10^5 cells per dish in 35-mm MatTek dishes with a 14-mm glass-bottom area (P35G-1.5-14-C, MatTek Corporation) for live-cell imaging. NucBlue (Thermo Fisher Scientific, Cat# R37605) was added before imaging to visualize nuclei. Images were acquired using a Leica TCS SP8 confocal laser-scanning microscope mounted on a Leica DMi8 microscope equipped with a motorized stage and a 63× oil-immersion objective. A white-light laser was used for excitation, and images were acquired using Leica Application Suite X (LAS X). Acquisition settings were optimized to minimize spectral crosstalk between channels.

Image analysis was performed using Fiji/ImageJ (*72*). A custom Python script was used for three-dimensional segmentation and volume measurements. Image stacks were separated into nuclear and cellular channels. For cell segmentation, the cellular channel was duplicated and median and Gaussian filters (radius = 5 and σ = 3, respectively) were applied to the entire stack. A three-dimensional binary mask was generated using Li thresholding (*73*), followed by Distance Transform Watershed 3D segmentation (*74*) using Borgefors (3,4,5) distances, 16-bit output, normalization, dynamic = 4 and connectivity = 6. The resulting mask was analyzed using the 3D Objects Counter plugin (*75*) to determine cell volume and surface area. Within the segmented cell regions, the nuclear channel was duplicated and median and Gaussian filters (radius = 7 and σ = 3, respectively) were applied to the entire stack. Nuclear volume and surface area were then quantified using the 3D Objects Counter plugin (*75*) to measure the nuclear volume (*μ*m^3^) and surface (*μ*m^2^).

### Generation of RPL29–HaloTag knock-in cell lines using CRISPR–Cas9 gene editing

RPL29–HaloTag CRISPR knock-in vectors were used as described previously (*76*). HCT116 cells were co-transfected with donor and gRNA plasmids at a 1:1 ratio using Lipofectamine 2000 in Opti-MEM. Five days after transfection, cells were labelled with 100 nM HaloTag TMR ligand (Promega, Cat# G8251) to select positive clones. TMR-positive single cells were sorted using a MoFlo Astrios (Beckman Coulter Life Sciences) into 96-well plates containing RPMI supplemented with 33% FBS, 3.3% penicillin/streptomycin and 0.33% gentamicin. Single-cell clones were expanded and genetically screened for HaloTag insertion.

### Micro-patterning

Fibronectin-coated micropatterned disks (250 µm diameter) were fabricated using previously described photolithography methods (*77*). Glass coverslips were activated using a plasma cleaner (Harrick Plasma) and coated with PLL-g-PEG (Surface Solutions GmbH, 0.5 mg/mL in 10 mM HEPES). After washing with 1× PBS and deionized water, the surface was illuminated with deep UV light (UVO Cleaner, Jelight) through a chromium photomask (JD-Photodata). Coverslips were then incubated with fibronectin (Sigma-Aldrich, Cat# F1056; 20 µg/mL in 100 mM NaHCO₃, pH 8.4). Cells were detached using 0.02% EDTA (Versene, Gibco, Cat# E6758), seeded onto the micropatterned coverslips and allowed to attach for 16 h.

### R110 and TMR incorporation on micropatterns

Two-colour Ribo–Halo labelling was adapted from a previously described method (*76*). Micropatterned HCT116 RPL29–Halo cells were grown to confluence and incubated with 50 nM HaloTag R110 ligand for 2 h to label pre-existing RPL29. Cells were then washed three times with 2 mL fresh RPMI for 10 min each in the dark to remove unbound R110 ligand. Following washing, cells were either maintained under control conditions or subjected to 10 kPa compression for 30 min in RPMI containing 100 nM HaloTag TMR ligand to label newly synthesized RPL29. Cells were subsequently washed as above and fixed in 4% paraformaldehyde for 10 min at room temperature. After fixation, cells were washed, permeabilized, stained with DAPI and mounted for imaging. Confocal imaging was performed using a Leica TCS SP5 confocal laser-scanning microscope mounted on a Leica DMI 6000B inverted microscope and equipped with an HCX PL APO 63×/1.4 NA oil-immersion objective. Images were acquired using Leica LAS AF software with 405-, 488– and 561-nm laser lines.

### Fluo-8® AM Imaging

HCT116 cells were seeded at 4 × 10^5 cells per dish in 35-mm MatTek dishes with a 14-mm glass-bottom area (P35G-1.5-14-C, MatTek Corporation). After 24 h, culture medium was replaced with calcium-imaging buffer containing 20 mM HEPES, 135 mM NaCl, 5 mM KCl, 0.4 mM KH₂PO₄, 1 mM MgSO₄, 5.5 mM glucose and 1 mM Ca²⁺. Cells were loaded with 5 µM Fluo-8 AM (Abcam, Cat# ab142773) for 30 min before imaging. Where indicated, cells were pre-treated with 3 µM thapsigargin (Selleckchem, Cat# S7895) for 5 min, followed by drug washout and a 30-min recovery period before imaging. Compression was applied using 10-kPa polyacrylamide gels at the start of image acquisition. NucBlue (Thermo Fisher Scientific, Cat# R37605) was added before imaging to visualize nuclei.

Live-cell imaging was performed using a Leica TCS SP8 confocal laser-scanning microscope mounted on a Leica DMi8 microscope equipped with a motorized stage and a 63× oil-immersion objective. A white-light laser was used for excitation, and images were acquired using Leica Application Suite X (LAS X). Baseline fluorescence was recorded before compression, and Fluo-8 fluorescence was monitored continuously for 30 min after compression onset. Image analysis was performed using Fiji/ImageJ.

### EdU incorporation

Micropatterned HCT116 cells were grown to confluence, and EdU incorporation was assessed using the Click-iT™ Plus EdU Cell Proliferation Kit for Imaging with Alexa Fluor™ 488 dye (Thermo Fisher Scientific, Cat# C10367). Cells were incubated with 10 µM EdU and subsequently processed according to the manufacturer’s instructions (https://documents.thermofisher.com/TFS-Assets/LSG/manuals/mp10637.pdf).

Confocal imaging was performed using a Leica TCS SP5 confocal laser-scanning microscope mounted on a Leica DMI 6000B inverted microscope and equipped with an HCX PL APO 63×/1.4 NA oil-immersion objective. Images were acquired using Leica LAS AF software with 405– and 488-nm laser lines.

### p-Body quantification in fixed cells

Cells were seeded at approximately 30,000 cells/cm² onto 13-mm glass coverslips pre-coated for 2 h with 10 µg/mL fibronectin from human plasma (Sigma-Aldrich, Cat# F1056). After 24 h, cells were maintained in complete medium and subjected to 10 kPa compression for 2.5, 5, 10 or 15 min. Parallel cultures that were not compressed were used as controls. Following mechanical stress, cells were fixed in 4% paraformaldehyde (HiMedia, Cat# TCL119-100ML) for 10 min and washed three times with 1× DPBS. Cells were permeabilized with 1× PBS containing 0.1% Triton X-100 for 5 min at room temperature and washed three times with 1× PBS. Blocking was performed in 1× PBS containing 5% normal donkey serum (Jackson ImmunoResearch, Cat# 017-000-121). Cells were incubated overnight at 4 °C with anti-DCP1A antibody (Abcam, Cat# ab47811) diluted 1:800 in blocking solution. After three washes of at least 10 min each with 1× PBS, cells were incubated for 1 h at room temperature with Alexa Fluor 647 donkey anti-rabbit secondary antibody (Thermo Fisher Scientific, Cat# A-31573) diluted 1:400 together with TRITC-phalloidin (Sigma-Aldrich, Cat# P1951) diluted 1:50. Cells were washed three times with 1× PBS and stained with DAPI (Sigma-Aldrich, Cat# D9542) diluted 1:5,000 for 15 min, followed by three additional washes. Coverslips were mounted in a medium composed of 70% glycerol, 30% 1× PBS and 1.5% DABCO.

Confocal microscopy was performed using a Leica TCS SP5 confocal laser-scanning microscope based on a Leica DMI 6000B inverted microscope. Images were acquired using an HCX PL APO 63×/1.4 NA oil-immersion objective and Leica LAS AF software. Laser lines of 405, 488, 561 and 633 nm were used. Images were acquired as z-stacks with a 0.3-µm step size at 2048 × 2048 resolution. Approximately 30 cells per condition and per replicate were analyzed using the custom Fiji workflow described below.

Cell bodies were segmented from phalloidin staining as follows. Background was first subtracted using a rolling-ball radius of 200 pixels, followed by Gaussian and variance filtering (σ = 3 and radius = 7, respectively). Segmentation was then performed using the MinError thresholding method in Fiji. To improve object detection, segmented images were subjected to closing, hole filling and five erosion steps before application of the Analyze Particles plugin (100 µm² < size < 800 µm²) to identify cell bodies. Cell doublets were manually removed. P-bodies were segmented after background subtraction using a rolling-ball radius of 4 pixels and thresholding at 25. For each cell-body ROI, the Analyze Particles plugin (size < 10 µm²) was used to identify granules. For each cell, the following parameters were extracted: cell area, granule count, mean granule area, minimum granule area, maximum granule area and standard deviation of granule area.

### 2′,7′-dichlorofluorescin diacetate (DCFDA) flow cytometry

Cells were seeded at approximately 3 × 10^5^ cells per well in 6-well plates (Falcon, Cat# 353046) 1 d before treatment. Cells were subjected to 10 kPa compression using an elastic overlay. After approximately 10 min, cells were harvested by trypsinization and pelleted at 1200 rpm for 5 min. Pellets were resuspended in 1× DPBS supplemented with 1% dialyzed fetal bovine serum (EuroClone, Cat# ECS0181L) and 20 μM DCFDA (Sigma-Aldrich, Cat# D6883). Approximately 10 min after resuspension, samples were analyzed by flow cytometry on an Attune NxT instrument (Thermo Fisher Scientific) equipped with a 488-nm laser. Mean BL1-A values were recorded for each sample. Data were analyzed using FlowJo software.

### Flow cytometry–based analysis of mitochondrial functionality

Cells were seeded at 7 × 10^5 cells per well in 6-well plates 1 d before treatment. Before compression, cells were washed twice with PBS to remove serum and pre-incubated with 200 nM MitoTracker Green (Thermo Fisher Scientific, Cat# M7514) for 30 min to label mitochondrial mass. Cells were then incubated with 20 nM TMRM (Thermo Fisher Scientific, Cat# T668) and either maintained under control conditions or subjected to 30 min of compressive stress. MitoTracker Green was maintained during the compression step, and compression gels were pre-equilibrated with medium containing 20 nM TMRM before use. At the end of treatment, cells were washed with PBS, detached rapidly using a minimal volume of trypsin and collected in 1 mL PBS. Cell suspensions were transferred to 1.5-mL tubes, pelleted by centrifugation, resuspended in PBS and transferred to FACS tubes for acquisition. Control samples included unstained cells, MitoTracker Green only, TMRM only, MitoTracker Green + TMRM, and the corresponding compressed conditions. DAPI dilactate (Sigma-Aldrich, Cat# D9564; stock 1 mg/mL) was added immediately before acquisition to exclude dead cells (5 µL per sample). Samples were acquired on a 5-laser Aurora cytometer (Aurora 5L, Cytek Biosciences, USA) and analyzed using FlowJo software (v10.10, BD Biosciences, USA). Mitochondrial functionality was defined as the ratio between mitochondrial membrane potential and mitochondrial mass and was calculated in live cells as the ratio between TMRM median fluorescence intensity and MitoTracker Green median fluorescence intensity.

### TMR incorporation for in-gel fluorescence analysis of Ribo–Halo degradation

Ribo–Halo labelling was adapted from a previously described method (*76*). HCT116 RPL29–Halo cells were plated in 6-well plates at 7 × 10^5 cells per well. After 24 h, cells were incubated with 100 nM HaloTag TMR ligand for 1 h to label pre-existing ribosomes. Cells were then washed three times and incubated in RPMI for 0.5, 1, 2 or 3 h, either under control conditions or subjected to 10 kPa compression. Cells were subsequently trypsinized, lysed in RIPA buffer and sonicated three times at 4 °C. Following Bradford protein quantification, 15 µg of lysate was resolved by SDS–PAGE on 4–12% Bolt gels (Thermo Fisher Scientific). In-gel TMR fluorescence was detected using an iBright fluorescence imaging system (Thermo Fisher Scientific). Gels were subsequently stained with Coomassie blue as a loading control, and TMR fluorescence was normalized to total protein and quantified using Fiji.

### Azidohomoalanine (AHA) incorporation for in-gel fluorescence analysis of protein degradation

Protein degradation kinetics were assessed by adapting a previously published protocol (*76*). Briefly, equal numbers of HCT116 cells were plated in 6-well plates 24 h before the experiment. Cells were washed three times with PBS, and the medium was replaced with methionine-free RPMI (Thermo Fisher Scientific, Cat# A1451701) supplemented with either 250 µM methionine or 250 µM AHA. Cells were incubated for 3 h to allow AHA incorporation and then washed three times with RPMI (Thermo Fisher Scientific, Cat# 61870036). Cells were either maintained under control conditions or subjected to 10 kPa compression for the indicated time course. Cells were then washed three times with PBS, counted, and equal cell numbers were lysed in RIPA buffer followed by sonication. Samples were reduced with 5 mM TCEP and subsequently alkylated with chloroacetamide at a final concentration of 20 mM. Lysates were then subjected to click-chemistry reactions containing final concentrations of 1 mM CuSO₄, 1 mM sodium ascorbate, 100 µM TBTA ligand and 100 µM TMR-alkyne. The click-reaction mixture was incubated for 1 h in the dark at room temperature. Samples were resolved by SDS–PAGE on 4–12% Bolt gels (Thermo Fisher Scientific). In-gel TMR-alkyne fluorescence was detected using an iBright fluorescence imaging system (Thermo Fisher Scientific). Gels were subsequently stained with Coomassie blue as a loading control, and TMR fluorescence was normalized to total protein and quantified using Fiji/ImageJ.

### Redox western blot

Redox-dependent cysteine modifications were analyzed by non-reducing PEG-switch western blotting. Cells were lysed in a non-reducing HEPES-based buffer containing 50 mM HEPES, pH 7.4, 150 mM NaCl, 1% Triton X-100, 1 mM EDTA and protease inhibitor cocktail (Calbiochem, Cat# 539138), and protein concentration was determined using a BCA assay (Thermo Fisher Scientific, Cat# 23225). Control lysates were divided into three aliquots: (A) direct labeling with mPEG2K-maleimide (Cat# JKA3124) to detect reduced cysteines present in vivo; (B) reduction with TCEP (Thermo Fisher Scientific, Cat# 20490) followed by mPEG labeling to generate a fully reduced control; and (C) reduction followed by alkylation with N-ethylmaleimide (NEM; Sigma-Aldrich, Cat# E1271) to block free thiols and prevent PEG labeling. Experimental samples, including control, compressed, NAC-treated and H₂O₂-treated conditions, were processed according to condition A. H₂O₂ was used at 150 µM as an acute oxidation-positive control and applied for the same duration as the corresponding compression treatment.

For mPEG labeling, samples were incubated under non-reducing conditions with gentle mixing. Following labeling, proteins were precipitated with 80% cold acetone. For 100 µL lysate, 440 µL acetone was added; for 150 µL lysate, 600 µL acetone was added. Samples were centrifuged at 15,000 × g for 10–15 min at 4 °C, supernatants were carefully removed and pellets were air-dried for 1–2 min. Pellets were resuspended in non-reducing Laemmli buffer and heated at 65–70 °C for 5 min before electrophoresis.

Proteins were separated by non-reducing SDS–PAGE and transferred to nitrocellulose membranes. Membranes were probed with antibodies against eIF2β, eIF5 and eEF2. PEG-dependent electrophoretic mobility shifts were interpreted as indicators of cysteine oxidation state.

### Electron microscopy

Correlative light-electron microscopy (CLEM), conventional electron microscopy and electron tomography were performed as previously described (*78*). A brief description of the procedures is provided below.

#### Embedding

Cells were grown on MatTek glass-bottom dishes (MatTek Corporation, USA) and fixed in a mixture of 2.5% paraformaldehyde and 2.5% glutaraldehyde (EMS, USA) in 0.1 M sodium cacodylate buffer, pH 7.4, for 2 h at room temperature. Samples were then washed six times in 0.2 M sodium cacodylate buffer, pH 7.2, and incubated for 1 h at room temperature in a 1:1 mixture of 2% osmium tetroxide (OsO₄) and 3% potassium ferrocyanide. Samples were subsequently washed six times in cacodylate buffer, pH 6.9, and treated sequentially with 0.3% thiocarbohydrazide in 0.2 M cacodylate buffer for 10 min and 1% OsO₄ in 0.2 M cacodylate buffer, pH 6.9, for 30 min. Samples were then washed with 0.1 M sodium cacodylate buffer, pH 6.9, until all traces of osmium fixative were removed, rinsed in deionized water, incubated with 1% uranyl acetate in water for 1 h and washed again in water. Samples were dehydrated through an ethanol series, embedded in epoxy resin at room temperature and polymerized at 60 °C for at least 72 h.

#### Sectioning

Cells of interest were selected by confocal imaging of MatTek dishes using optical sectioning and Z-stack acquisition. Embedded samples were sectioned using a diamond knife (Diatome, Switzerland) on a Leica EM UC7 ultramicrotome (Leica Microsystems, Vienna). Serial 70-nm sections were collected for conventional electron microscopy, whereas 200-nm semi-thick sections were collected for electron tomography. Sections were examined using a Tecnai20 transmission electron microscope (FEI, now Thermo Fisher Scientific, The Netherlands) operated at 200 kV.

#### Electron tomography

Serial 200-nm semi-thick sections were collected on 1% Formvar films mounted on slot grids. Both sides of the grids were labelled with 10-nm fiducial gold particles (PAG10, CMC, Utrecht, The Netherlands). Tilt series were acquired from ±65° at 1° increments at 200 kV using a Tecnai20 electron microscope (FEI, now Thermo Fisher Scientific, Eindhoven, The Netherlands). Tilt series were acquired at magnifications of 9,600×, 14,500×, 19,000× or 29,000× using software supplied with the instrument. The nominal resolution of the tomograms was approximately 3 nm, based on section thickness, number of tilts, tilt increment and angular range. Individual tomograms were reconstructed using the IMOD package, and membrane contours were assigned using 3DMOD v4.0.11.

## Imaging and Single-particle tracking

### Image acquisition highly inclined and laminated optical sheet (HILO)

Movies of pCMV-PfV-Sapphire-IRES-DSRed-infected HCT116 cells were acquired on a custom-built HILO microscope capable of single-molecule imaging (*79*). The system comprised an illumination arm connected to a commercial microscope frame (Olympus IX-71, Evident). The illumination arm directed light from a 200 mW 561-nm laser (Cobolt 06-DPL, Hübner Photonics) to perform single-molecule tracking using HILO illumination. A movable mirror in a plane conjugated to the back focal aperture of the objective allowed tuning of the excitation angle in the object plane.

Imaging was performed with a 60× 1.49 NA oil-immersion objective (Olympus ApoN 60× 1.49 Oil, Evident). Fluorescence emitted from the sample was collected through the same objective, filtered with a quad-band emission filter (FF01-466/523/600/677-25, Semrock) and detected using an sCMOS camera (Orca Fusion C14440-20UP, Hamamatsu Photonics). Videos were acquired at 100 frames per second using Micro-Manager. The microscope was equipped with Okolab environmental control systems for temperature (37 °C), CO₂ (5%) and humidity to maintain cells under physiological conditions during live-cell imaging.

Particle tracking was performed using the MosaicSuite particle-tracking plugin in Fiji/ImageJ, assuming Brownian dynamics. Typical tracking parameters were a particle radius of 3 pixels, an intensity cut-off of 0.001, a link range of 1 frame and a maximum displacement of 5 pixels.

### Extraction of the rheological parameters

For every trajectory, we calculated the time-averaged mean square displacement (MSD) as defined in (*25*, *80*). To characterize individual particle trajectories, we calculated apparent diffusion coefficients by fitting the MSD with a linear (diffusive) time dependence at short time scales, as in (*25*).

### Phospho-Proteomics

#### Cell lysis

Cells were seeded at approximately 3 × 10^5 cells per well in 6-well plates (Falcon, Cat# 353046) 24 h before harvesting. On the day of the experiment, cells were subjected to 10 kPa compression using an elastic overlay. Following compression, cells were washed once with PBS and lysed directly in the well using 8 M urea in 100 mM Tris-HCl, pH 8.0. Lysates were collected, sonicated for three cycles of 30 s on/30 s off at 14 °C using a Bioruptor, and cleared by centrifugation at 12,000 rpm for 20 min. Cleared supernatants were collected and protein concentrations were determined using the Pierce BCA Protein Assay (Thermo Fisher Scientific, Cat# 23225).

### Protein digestion and peptide purification

For each sample, 200 µg of protein from 8 M urea lysates was digested in solution. Disulfide bonds were reduced with TCEP to a final concentration of 10 mM and cysteines were alkylated with 2-chloroacetamide to a final concentration of 40 mM. Samples were incubated for 30 min at room temperature. Samples were diluted with 100 mM Tris-HCl, pH 8.0, to 5 M urea and digested with Lys-C at an enzyme:protein ratio of 1:50 for 1 h at room temperature. Samples were then further diluted to 2 M urea with 100 mM Tris-HCl, pH 8.0, and digested overnight at room temperature with trypsin at an enzyme:protein ratio of 1:44 (Kulak et al. 2014).

After acidification to 5% formic acid, peptides were desalted using Sep-Pak C18 cartridges (C18 Classic Cartridge, Waters). Cartridges were conditioned with 1 mL 50% acetonitrile in 5% formic acid and 1 mL 5% formic acid, followed by sample loading and washing with 1 mL 5% formic acid. Peptides were eluted with two 500-µL volumes of 80% acetonitrile in 5% formic acid. Acetonitrile was removed in a SpeedVac, and samples were resuspended in 50 µL 0.2% trifluoroacetic acid for phosphopeptide enrichment.

### Phosphopeptide enrichment

Samples were processed using a sequential phosphopeptide-enrichment strategy combining Ti-IMAC magnetic beads (MagReSyn Ti-IMAC, Resyn Biosciences) and TiO₂ beads (Titansphere 5 µm, GL Sciences). For each sample, 40 µL Ti-IMAC HP magnetic beads (20 µg/µL) were equilibrated with 70% ethanol, elution buffer 1 (1% NH₄OH) and loading buffer (80% acetonitrile, 5% TFA, 1 M glycolic acid), and incubated with the samples for 20 min with mixing. Beads were washed with wash buffer 1 (80% acetonitrile, 1% TFA) and wash buffer 2 (10% acetonitrile, 0.2% TFA), and phosphopeptides were eluted with elution buffer 1 and elution buffer 2 (50% acetonitrile, 5% NH₄OH).

The Ti-IMAC flow-through was subjected to a second phosphopeptide-enrichment step using TiO₂ beads. For each sample, 4 mg TiO₂ beads were loaded into a tip containing a C8 plug and equilibrated with buffer 1 (80% acetonitrile, 0.2% TFA, 1 M glycolic acid). The Ti-IMAC flow-through was loaded onto the beads after centrifugation at 1,500 × g for 5 min. Beads were washed with buffer 1 and buffer 2 (80% acetonitrile, 0.2% TFA). Phosphopeptides were eluted with buffer 3 (1% NH₄OH) and buffer 4 (60% acetonitrile). Pooled Ti-IMAC and TiO₂ eluates were dried in a SpeedVac and resuspended in 12 µL 5% formic acid for LC–MS/MS analysis.

### nLC and mass spectrometry

Phosphoproteomic analysis was performed on a Vanquish Neo nanoLC coupled to a Quadrupole Orbitrap Exploris 480 mass spectrometer equipped with a Nano Flex ESI source and the FAIMS Pro interface (Thermo Fisher Scientific, Bremen, Germany). Phosphopeptides were separated on a 25-cm, 75-µm inner-diameter nano-column packed in-house with ReproSil-Pur C18-AQ 1.9-µm beads (Dr. Maisch) at a flow rate of 200 nL/min using a 131-min gradient. Each sample was analyzed in technical replicate using FAIMS compensation voltages of −30/−50 V and −40/−60 V in positive data-independent acquisition mode with HCD fragmentation. Full MS spectra (400–1,000 Th) were acquired at 120,000 resolution with a normalized AGC target of 300%, a maximum injection time of 55 ms and one microscan. Fragment spectra were acquired at 30,000 resolution with a normalized AGC target of 1,000%, automatic maximum injection time, one microscan, normalized collision energy of 32% and an isolation window of 30 m/z.

### Data processing and analysis

Raw data were processed in Spectronaut (v18.5) using a directDIA+ search with an in-house phosphopeptide library extension. Trypsin was specified as the cleavage enzyme with up to two missed cleavages. Carbamidomethylation of cysteine was set as a fixed modification, whereas methionine oxidation, protein N-terminal acetylation and phosphorylation of serine, threonine and tyrosine were set as variable modifications. A false-discovery rate of 0.01 was applied at both peptide and protein-group level. PTM localization filtering was enabled using a localization probability cutoff of 0.75. Quantification was performed using MS1 intensity, and cross-run normalization was disabled. Statistical analysis was performed in Perseus (v2.0.10.0) after quantile-centering normalization of phosphopeptide intensities. Volcano plots were generated in R (v4.3.1).

### Lysosome purification

HCT116 cells were transduced with lentivirus carrying pLJM1-Tmem192-mRFP-3xHA (Addgene plasmid #134631) and selected with 1 µg/mL puromycin for 5 d to generate a stable cell population for lysosome immunopurification experiments. Immunoprecipitation was performed as previously described (*81*). Briefly, approximately 35 million cells per condition (control and 1 h 10 kPa compressed) were harvested in KPBS (136 mM KCl, 10 mM KH₂PO₄, pH 7.25 adjusted with KOH). Following centrifugation, the pellet was resuspended in 1 mL KPBS and subjected to 20 strokes of Dounce homogenization. A 25-µL aliquot was then set aside for the whole-cell fraction; the remaining lysate was centrifuged at 1,000 × g for 2 min at 4 °C, and the supernatant was incubated with 150 µL of KPBS-prewashed anti-HA magnetic beads on a gentle rotator for 3 min. Immunoprecipitates were washed gently three times in KPBS and eluted using 8 M urea in 100 mM Tris-HCl, pH 8.0. The whole-cell fraction was diluted in 10 volumes of 8 M urea in 100 mM Tris-HCl, pH 8.0.

### Proteomic Analysis of Lysosome-Enriched Fractions and Whole Cell Extracts Sample Preparation and Protein Digestion

For lysosome-enriched samples, 10 µg of lysosomal enriched proteins were digested using the filter-aided sample preparation (FASP) protocol as described in (*82*). Lysosomal extracts were transferred to 10 kDa molecular weight cut-off Microcon filters (Millipore, Cat# MRCPRT010), and buffer exchange was performed with 8 M urea in 100 mM Tris-HCl, pH 8.0 (UA buffer). Reduction and alkylation were carried out in UA buffer supplemented with 10 mM TCEP (Thermo Scientific, cat. no. 20490) and 40 mM 2-chloroacetamide (Sigma-Aldrich, Cat# C0267) for 30 min at room temperature, followed by centrifugation at 10,000 × g for 15 min. Filters were washed twice with 100 µL UA buffer and twice with 100 µL 100 mM Tris-HCl, pH 8.0. Proteins were digested by adding 0.2 µg Lys-C in 95 µL 100 mM Tris-HCl, pH 8.0 and incubating at 37°C for 1 h. Subsequently, 0.3 µg trypsin was added, and samples were incubated overnight at 37°C. Peptides were recovered by centrifugation, purified using C18 StageTips (Proxeon Biosystems) according to (*83*), and eluted in 10 µL of 5% formic acid.

For whole cell extracts (WCE), cells were resuspended in UA buffer, and 10 µg of protein were subjected to in-solution digestion. Reduction and alkylation were performed in 8 M urea, 100 mM Tris-HCl pH 8.0 containing 10 mM TCEP (Thermo Scientific) and 40 mM 2-chloroacetamide (Sigma-Aldrich) for 30 min at room temperature. Proteins were first digested with Lys-C at a 1:50 enzyme-to-protein ratio (w/w) in 100 mM Tris-HCl pH 8.0 containing 6 M urea for 1 h at 37°C. Samples were then diluted to 2 M urea and incubated overnight at 25°C with trypsin at a 1:30 enzyme-to-protein ratio (w/w). The resulting peptides were purified on C18 StageTips (Proxeon Biosystems) and eluted in 10 µL of 5% formic acid.

### LC–MS/MS Analysis

A total of 1.5 µL of purified peptides was injected into an Orbitrap Exploris 480 mass spectrometer (Thermo Fisher Scientific) equipped with a FAIMS Pro interface. Analytical columns were packed in-house with ReproSil-Pur C18-AQ resin (1.9 µm; Dr. Maisch GmbH) using a high-pressure bomb loader (Proxeon). Peptides were separated at a flow rate of 0.20 µL/min using a linear gradient from 95% solvent A (2% acetonitrile, 0.1% formic acid) to 30% solvent B (80% acetonitrile, 0.1% formic acid) over 62 min, followed by an increase from 30% to 60% solvent B over 5 min and from 60% to 100% solvent B over 13 min, resulting in a total run time of 80 min.

The mass spectrometer was operated in positive electrospray ionization mode using data-independent acquisition (DIA). MS1 scans were acquired at a resolution of 120,000 over a mass range of 400–1,000 m/z, with an automatic gain control (AGC) target of 3 × 10^6 and a maximum injection time of 55 ms. MS2 scans were acquired at a resolution of 30,000, with an AGC target of 1 × 10^7, maximum injection time set to Auto, higher-energy collisional dissociation (HCD) collision energy of 32% and 30 m/z isolation windows. The charge-state range was set from 2 to 6, the RF lens was set to 40% and loop control was set to “All”. The FAIMS Pro interface was operated with compensation voltages of −40 V and −60 V. The ion transfer tube temperature was set to 300 °C with a carrier gas flow of 3.7 L/min.

### DIA Data Processing and Database Search

Raw FAIMS-DIA files were analyzed using Spectronaut version 18 in DirectDIA mode with default BGS factory settings. Searches were performed against the UniProt canonical human proteome database, with trypsin specified as the digestion enzyme. Carbamidomethylation of cysteine was set as a fixed modification, whereas methionine oxidation and protein N-terminal acetylation were specified as variable modifications. False discovery rates were controlled at 1% at the peptide-spectrum match (PSM) and peptide/protein-group levels. Quantification was performed using the MS2 “Area” quantity type with cross-run normalization enabled.

## KPC eEF2 Knock-out

### Generation of eEF2K knockout Cell Line

sgRNAs targeting murine *Eef2k* were designed using vbc-score.org, with the *Mus musculus* mm10 genome as reference. The following sgRNAs were selected:

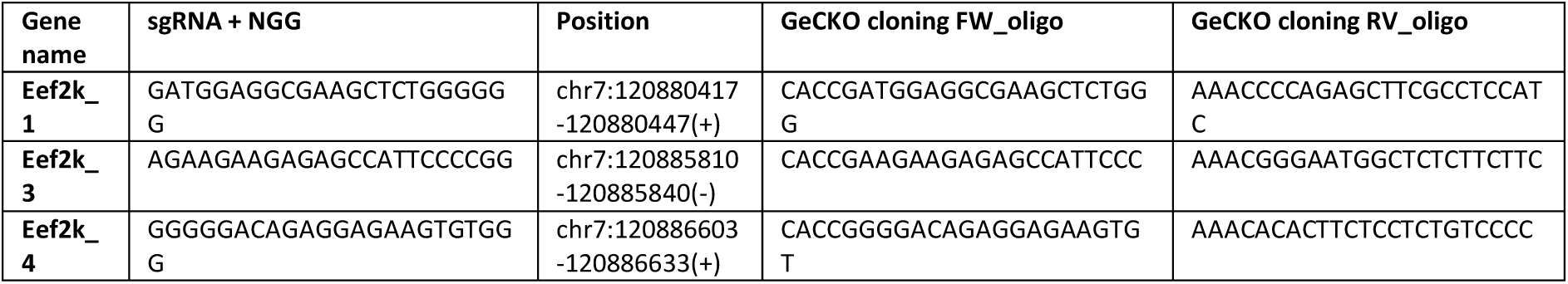

A guide targeting the Rosa safe-harbour locus was included as a control. Forward and reverse oligonucleotides for the Rosa-targeting guide (forward: CACCGCCACCCAGTGGAGTTAGTAT; reverse: AAACATACTAACTCCACTGGGTGGC) were obtained from LuBioScience GmbH. The annealed oligonucleotides were cloned into the lentiCRISPRv2 backbone (Addgene plasmid #52961) according to the Zhang laboratory Target Guide Sequence Cloning Protocol. Plasmids were transformed into NEB Stable Competent *E. coli* (New England Biolabs, Cat# C3040H), plated on LB-agar containing ampicillin and cultured overnight at 37 °C. Individual colonies were inoculated into 2 mL LB medium supplemented with 100 µg/mL ampicillin and cultured overnight at 37 °C with shaking at 200 rpm. Plasmid DNA was purified using the NucleoSpin Plasmid kit (Macherey-Nagel, Ref# 740588) and verified by Sanger sequencing at Microsynth AG.

For lentiviral production, HEK293T cells were seeded at 1 × 10^6 cells per well in 6-well plates containing 2 mL culture medium. Approximately 5 h later, cells were transfected with 2.9 µg vector, 1.3 µg psPAX2 and 0.7 µg VSV-G using Lipofectamine 3000 (Thermo Fisher Scientific, Cat# L3000008). The transfection mixture was incubated for 15 min at room temperature before addition to the cells, which were then incubated overnight at 37 °C and 5% CO₂. The following day, the medium was replaced with 2 mL fresh culture medium, and cells were incubated for a further 48 h.

Virus-containing supernatant was collected and filtered through a 0.45-µm filter before being added directly to KPC cells at approximately 50% confluence. Following overnight incubation, the viral medium was replaced with standard culture medium. Two days later, cells were selected in fresh culture medium supplemented with 10 µg/mL puromycin (InvivoGen, Cat# ant-pr-1). Selection was continued until all non-transduced control cells were eliminated, after which the transduced lines were returned to standard culture medium.

### Western blotting of KPC cells

KPC cells (CancerTools, Cat# 153474) were harvested in ice-cold RIPA lysis buffer containing 50 mM Tris-HCl (pH 7.5), 150 mM NaCl, 1% Triton X-100, 0.5% sodium deoxycholate and 0.1% SDS, supplemented with PhosSTOP phosphatase inhibitor (Roche, Cat# 04906837001). Lysates were incubated on ice for 15 min, sonicated three times for 10 s at 28% amplitude and clarified by centrifugation at 14,000 × g for 10 min at 4 °C. Protein concentration was determined using the Pierce BCA Protein Assay Kit (Thermo Fisher Scientific, Cat# 23225).

10 µg of protein was mixed with 6× loading buffer and denatured at 98 °C for 5 min. Protein samples were resolved on Bolt Bis-Tris Plus Mini Protein Gels (4–12%, 1.0 mm, WedgeWell; Thermo Fisher Scientific, Cat# NW04120BOX) for 1 h at 150 V using the Chameleon Duo Prestained Protein Ladder (LI-COR Biosciences, Cat# 928-60000). Proteins were transferred to Odyssey nitrocellulose membranes (LI-COR Biosciences, Cat# 926-31092) for 1 h at 10 V, and membranes were blocked with Intercept TBS Blocking Buffer (LI-COR Biosciences, Cat# 927-60001) for 30 min at room temperature.

Membranes were incubated overnight at 4 °C with anti-eEF2K antibody (Thermo Fisher Scientific, Cat# MA569976; 1:5,000) diluted in 2% BSA/TBS-T. After three washes in TBS-T, membranes were incubated for 30 min at room temperature with IRDye 800CW donkey anti-rabbit IgG secondary antibody (LI-COR Biosciences, Cat# 926-32213; 1:10,000) and imaged using an Odyssey CLx Imager (LI-COR Biosciences). Membranes were then washed three times in TBS-T and incubated for 1 h at room temperature with β-actin antibody (Santa Cruz Biotechnology, Cat# sc-47778; 1:2,000), followed by IRDye 680RD donkey anti-mouse IgG secondary antibody (LI-COR Biosciences, Cat# 926-68072; 1:10,000) for 30 min at room temperature. Blots were imaged again using the Odyssey CLx Imager.

### Animal experiments

All animal experiments were performed using 8-week-old male C57BL/6JRj mice obtained from Janvier. Animals were housed under specific-pathogen-free conditions in accredited animal facilities, with ad libitum access to food and water and a 12-h light/12-h dark cycle. KPC cells were collected at approximately 80% confluence using TrypLE Express (1×; Gibco, Cat# 12605-010). Following detachment, cells were collected in DMEM GlutaMAX (Gibco, Cat# 31966-021) supplemented with 10% fetal bovine serum and 5% penicillin/streptomycin, centrifuged at 290 × g for 5 min and resuspended in 1× PBS for cell counting and viability assessment. Mice were randomly assigned to experimental groups and orthotopically injected with 100,000 viable control or eEF2K-knockout KPC cells resuspended in 50 µL 1× PBS using a 28-gauge needle. Tumours were analysed 4 weeks after cell implantation. Animals were identified using coded IDs, and investigators performing tumour measurements were blinded to the experimental group.

At the experimental endpoint, mice were euthanized by gradually increasing the CO₂ concentration. Humane endpoints were defined as a palpable tumour exceeding 1.5 cm in diameter or body-weight loss exceeding 15%; these endpoints were not reached during the study. All procedures were performed in accordance with Swiss Federal regulations and were approved by the Cantonal Veterinary Office.

### Histology and immunohistochemistry

Haematoxylin and eosin staining (Diapath) was performed according to standard procedures, and sections were mounted using Eukitt mounting medium (Bio-Optica). Collagen deposition was assessed by Picrosirius Red staining (ScyTek Laboratories, SRS-IFU), and stained sections were digitized using a 3DHISTECH slide scanner (Epredia).

For immunohistochemistry, paraffin was removed with xylene and tissue sections were rehydrated through graded alcohol solutions. Antigen retrieval was performed for 30 min using preheated target-retrieval solution at pH 6. Sections were blocked for 60 min with FBS in PBS and incubated overnight with anti-phospho-eEF2 (Thr56) antibody (Cell Signaling Technology, Cat# 2331; 1:100). Antibody binding was detected using HRP-conjugated donkey anti-rabbit IgG (H+L), highly cross-adsorbed secondary antibody (Thermo Fisher Scientific, Cat# A16035) (1:500) for 2 h, followed by development with diaminobenzidine chromogen (DAB Peroxidase Substrate Kit, Vector Laboratories, Cat# SK-4100). Sections were counterstained with Mayer’s haematoxylin.

Immunohistochemical analyses were performed blinded by the histology facility. Samples were identified only by coded mouse IDs, and investigators performing the analyses were unaware of whether samples originated from control or eEF2K-knockout animals.

### Software and statistical analysis

Image analysis was performed using Fiji/ImageJ. Flow-cytometry data were analyzed using FlowJo (v10.10, BD Biosciences, USA). Statistical analyses and graph generation were performed using GraphPad Prism 10 for macOS (v10.6.1 (799), GraphPad Software), and figures were assembled using Adobe Illustrator (v30.1, Adobe). Proteomics data were processed using Spectronaut (v18.5), Perseus (v2.0.10.0) and R (v4.3.1).

Unless otherwise indicated, data are presented as mean ± SEM. Biological replicates represent independently performed experiments using separate cell cultures on different experimental days; technical replicates were not considered independent biological replicates. The number of independent experiments (*n*) and, where applicable, the number of individual cells or measurements analyzed are reported in the corresponding figure legends.

For comparisons between two matched conditions obtained within the same independent experiment, two-tailed paired Student’s *t*-tests were used, as indicated in the corresponding figure legends. Comparisons involving more than two matched conditions were performed using repeated-measures one-way ANOVA, whereas experiments involving multiple experimental conditions and time points were analyzed using two-way repeated-measures ANOVA. Mixed-effects models were used where repeated-measures datasets contained missing values. Independent groups were compared using unpaired two-tailed *t*-tests with Welch’s correction, where indicated. Multiple-comparisons corrections were applied as specified in the corresponding figure legends. Associations between normalized median GEM effective diffusion coefficients (*D*_eff_) and puromycin incorporation were assessed using Spearman rank correlation. Statistical significance was defined as *P* < 0.05. Exact statistical tests, sample sizes and significance thresholds are reported in the corresponding figure legends.

