## Supplementary Figures for "Compressive stress couples calcium signalling to translational control and proteostasis"

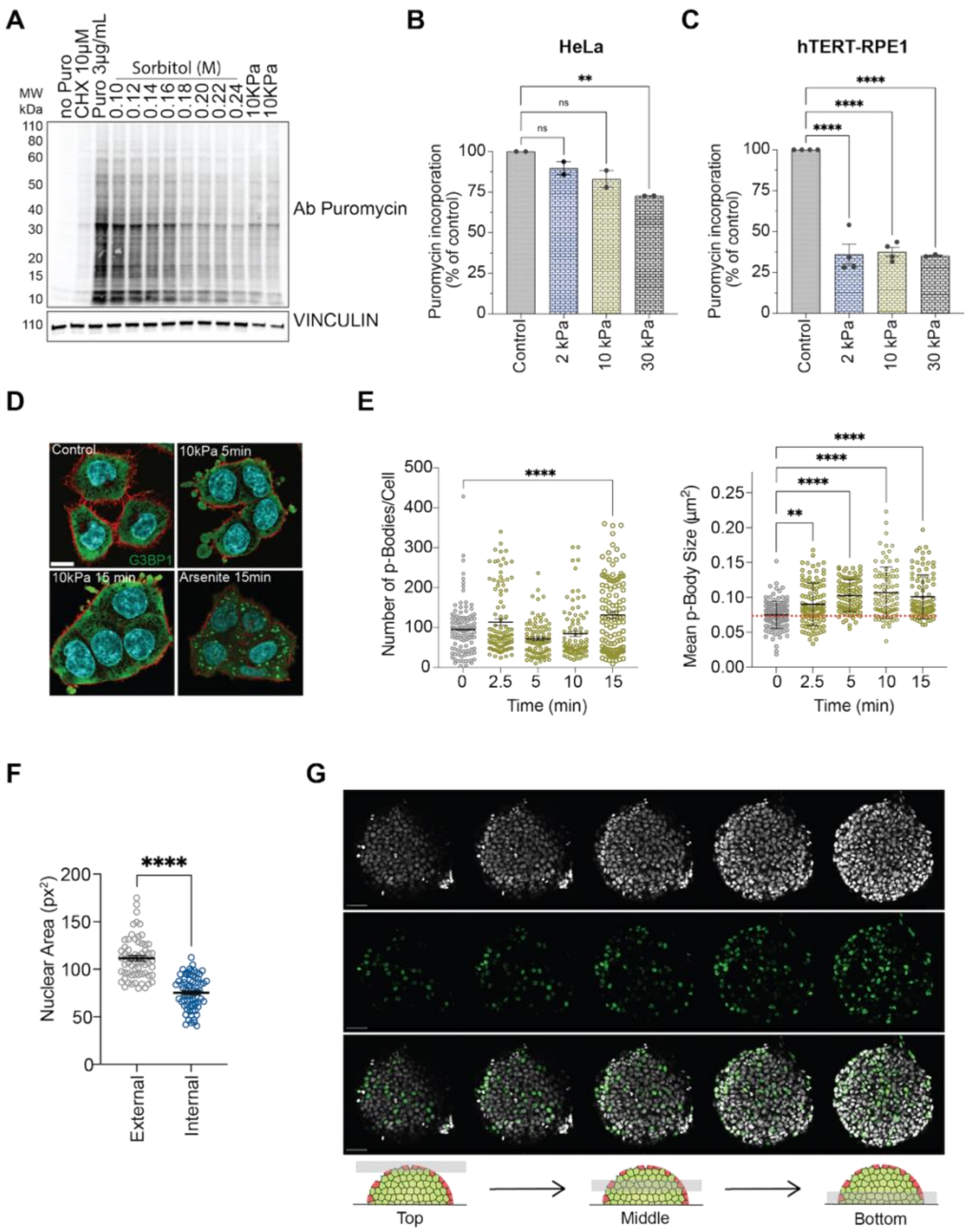


**Supplementary Figure 1. A.** Representative western blot of puromycin incorporation samples for HCT116 cells subjected to a range of osmotic (sorbitol) and 10 kPa compressive stress. Vinculin is used as a loading control. Data is representative of n=2 independent experiments. **B.** Puromycin incorporation in HeLa as measured by percentage of incorporation against control conditions for the indicated conditions. Points represent independent experiments, data is represented as Mean$\pm$SEM for n=2 independent experiments. **C.** Puromycin incorporation in hTERT RPE-1 as measured by percentage of incorporation against control conditions for the indicated conditions. Points represent independent experiments, data is represented as Mean$\pm$SEM n$\geq$3 independent experiments. **D.** DCP1A-positive P-body number and mean area per cell following 10 kPa compression for 0–15 min. Points represent individual cells pooled from three independent biological experiments; mean ± SEM. For control and compression at 2.5, 5, 10 and 15 min, respectively: n = 113, 106, 91, 87 and 117 cells for P-body number; n = 121, 106, 91, 87 and 93 cells for mean P-body area. **E.** Quantification of p-Bodies detected by immunofluorescence of Dcp1. Number and size of p-bodies were plotted following acute exposure to 10 kPa compression (0-15 minutes) were plotted and represented with Mean$\pm$SEM. Data is summed data from four independent replicates. **F.** Quantification of nuclear area of cells seeded and grown to confluency on 250 $\mu$m micropatterned disk patterns used in Fig. 1G and Supplementary Fig.1F) from 2 independent experiments. Data is represented as Mean$\pm$SEM. **G**. Representative EdU imaging of HCT116 cells grown on 250 $\mu$m micropatterned disks. The images from left to right represent planes taken from the top middle and bottom of the patterned cells. The scale bar represents 50 $\mu$m.

One way ANOVA with Sidak's multiple comparisons test (A,B,D). Unpaired t-test with Welsh's correction (E). $P$ values: $*$ $P < 0.05$, $**$ $P < 0.01$, $***$ $P < 0.001$, $****$ $P < 0.0001$.

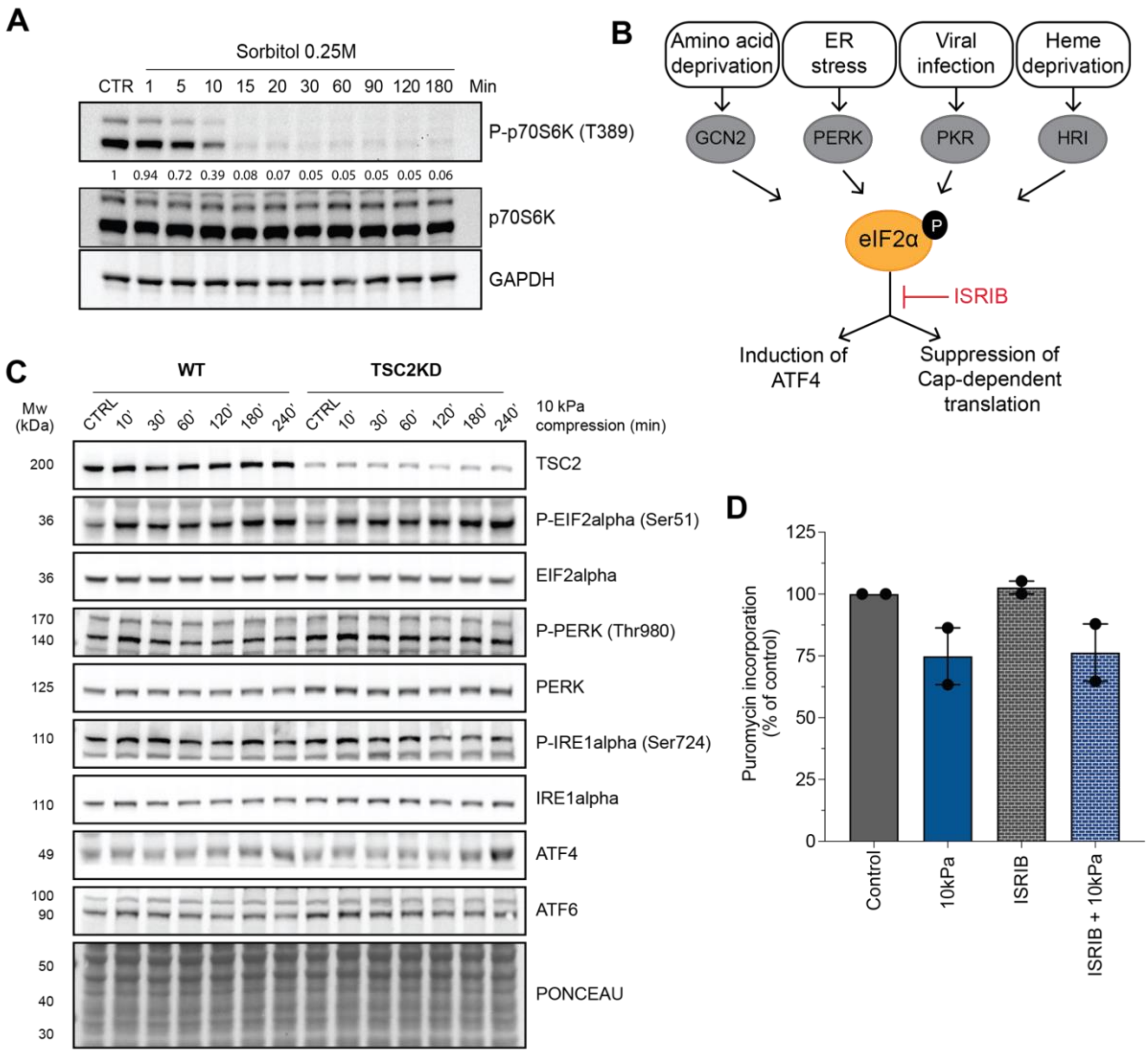


**Supplementary Figure 2. A.** Representative westerns showing kinetics of loss of P-p70S6K (T389) phosphorylation upon exposure to 0.25M Sorbitol. GAPDH was used as the loading control. Blots are representative of n=2 independent experiments. **B.** Schematic representation of stimuli and kinases involved in the integrated stress response, and the placement of ISR inhibitor ISRIB within the cascade. **C.** Representative western of activation of ISR associated kinases and downstream targets, eIF2, PERK, IRE1, ATF4 and ATF6 in control and TSC2KD cells in a time course spanning 0-6 hours of 10kPa compression. Ponceau S staining was used as a loading control. Blots are representative of n=2 independent experiments**.** Puromycin incorporation in control and 10kPa treated cells for DMSO and ISRIB (220 nM) treated conditions. Data points represent n=2 independent experiments, plotted as Mean ± SEM.

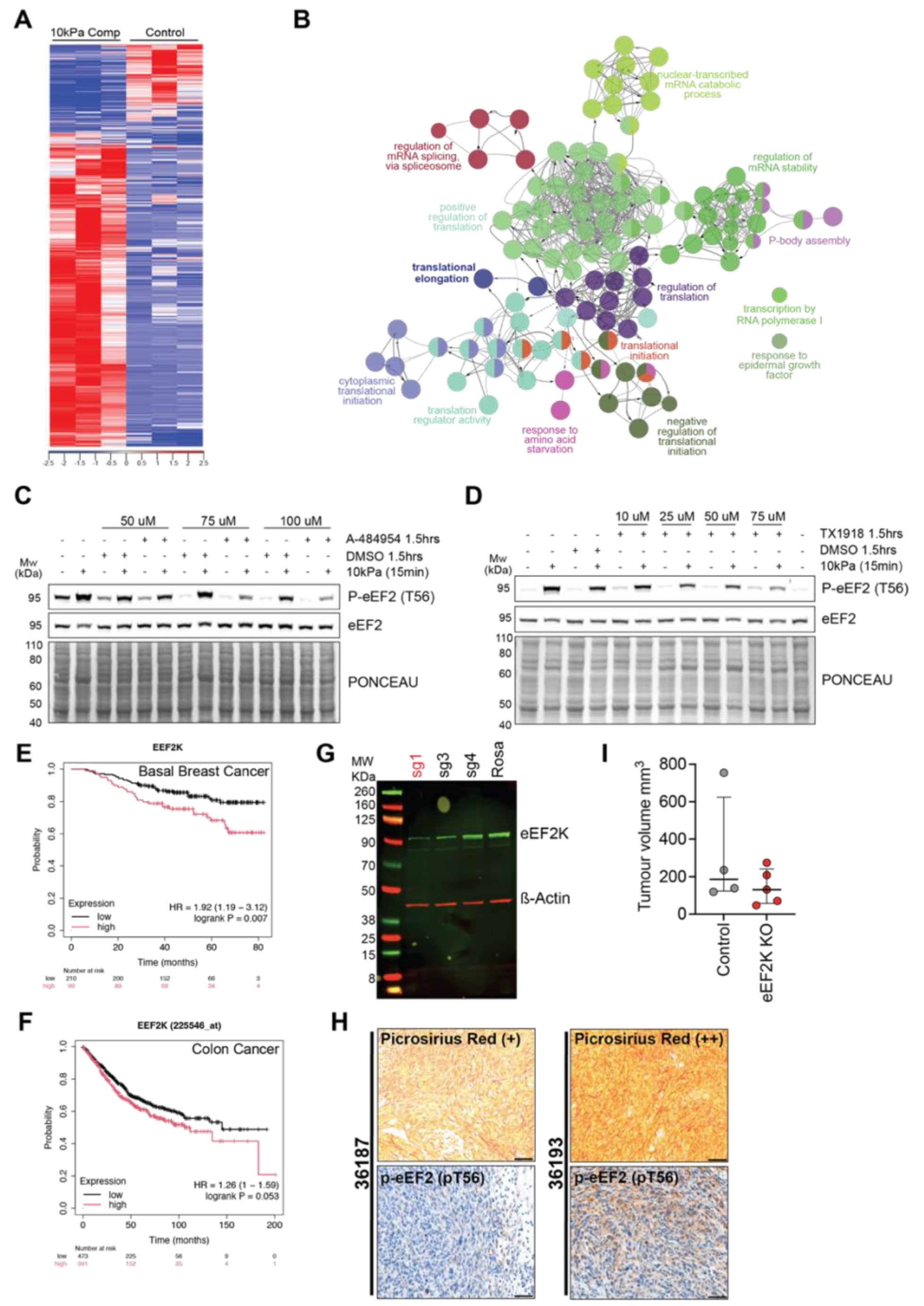

**Supplementary Figure 3. A.** Heatmap representing differentially phosphorylated peptides for control and 10 kPa compressed conditions. **B.** Translation-associated proteins with increased phosphorylation under compression were analysed using ClueGO in Cytoscape to identify the biological processes represented among the regulated proteins. Nodes represent enriched GO Biological Process terms, with related terms grouped according to functional similarity. Node size represents the number of associated proteins, and node colour denotes functional groups. **C.** Representative western of p-eEF2 (T56) under increasing concentrations of eEF2K inhibitor A-484954 in control and compressed conditions. Ponceau S staining serves as a loading control. Blots are representative of n=2 independent experiments. **D.** Representative western of p-eEF2 (T56) under increasing concentrations of eEF2K inhibitor TX1918 in control and compressed conditions. Ponceau S staining serves as a loading control. Blots are representative of n=2 independent experiments. **E.** Kaplan Meier plot of overall survival for high versus low eEF2 kinase expressing Basal Breast cancer patients. **F.** Kaplan Meier plot of overall survival for high versus low eEF2 kinase expressing Colon cancer patients **G.** Western blot of eEF2K levels in KPC control and KPC_eEF2K cell lines generated using the indicated synthetic guide RNAs. **H.** Representative Picrosirius Red and IHC for p-eEF2 (pT56) on two primary tumours (36187 and 36183) harvested following orthotopic injection of the control KPC-Rosa line**.** Scale bar represents 50$\mu$m. **I.** Tumour volumes were calculated using the ellipsoid formula (V = π/6 × L × W × H). Data are shown as individual data points with the median and interquartile range (IQR) indicated by black lines. Rosa, n = 4; KO, n = 6. *P*=0.4127 Mann Whitney test.

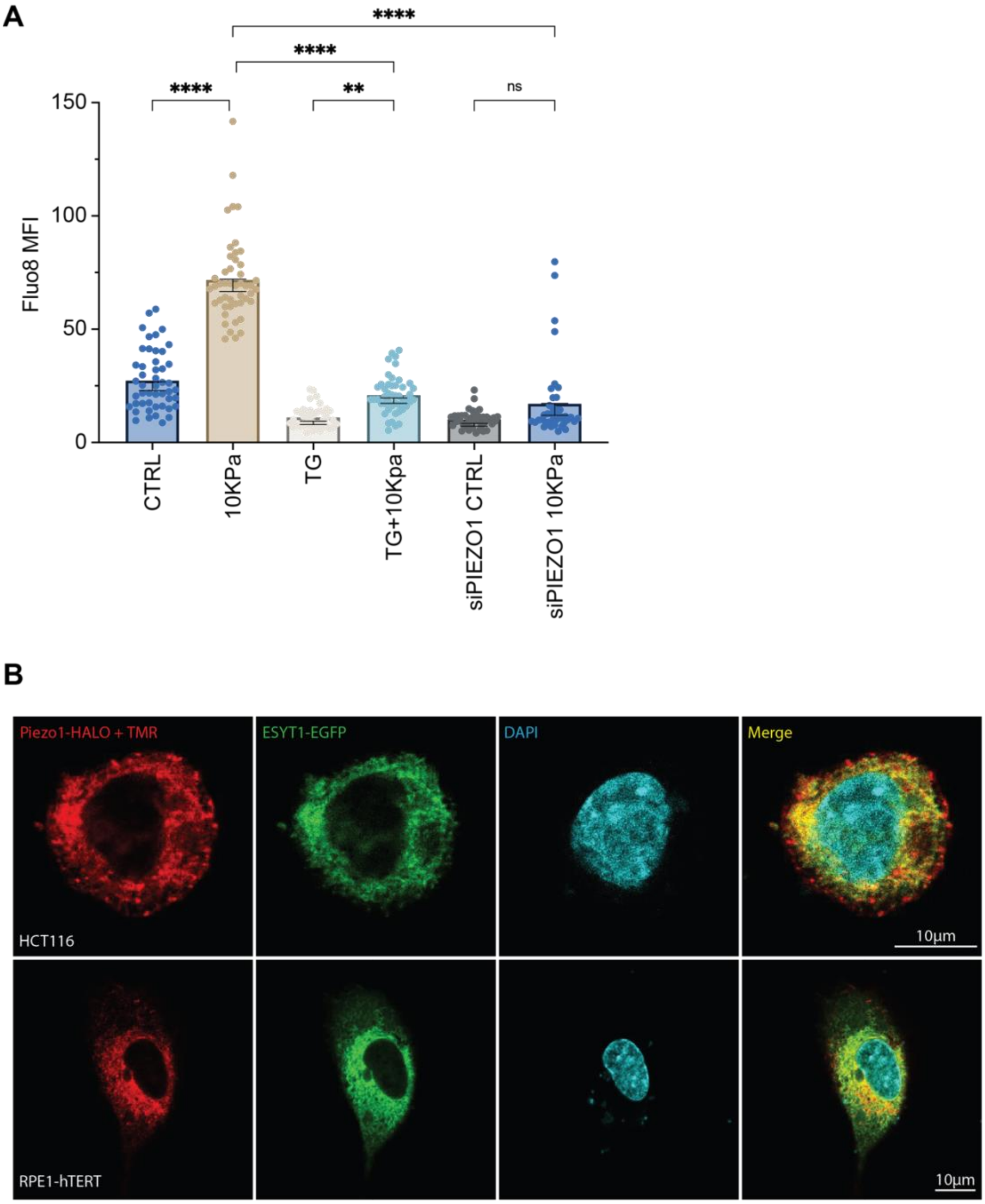


**Supplementary Figure 4. A.** Quantification of MFI for Fluo8 for the indicated conditions. Cells were treated with 2$\mu$M Tg for 5 minutes followed by a 30 minute washout before the beginning of imaging. Data points represent individual cells plotted as Mean $\pm$ SEM for pooled data from n=2 independent experiments **B.** Representative confocal microscopy images from HCT116 and hTERT-RPE1 cells co-transfected with Piezo1-Halo and ESYT1-EGFP constructs demonstrating Piezo1 localization to the ER. Piezo1 was visualized through addition of Halo ligand TMR. Scale bars represent 10 μm.

One-way ANOVA with Sidak's multiple comparison test(A). *P* values: ** $P < 0.01$, **** $P < 0.0001$.

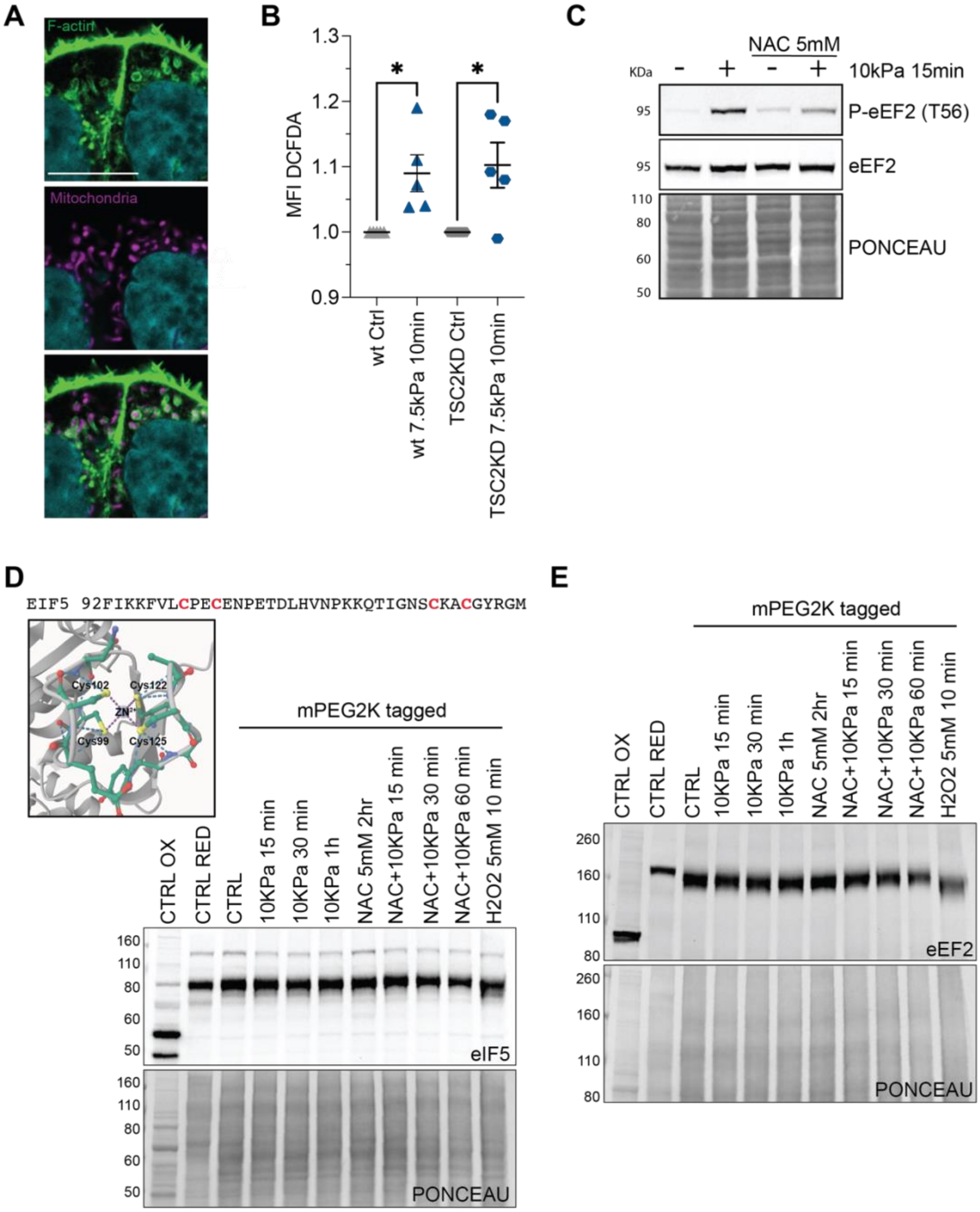


**Supplementary Figure 5. A.** Confocal imaging of Mito-dsRed with Phalloidin staining of F-Actin in 10 kPa compressed cells. Scale bar represents 10 $\mu$m. **B.** MFI of ROS indicator Dichlorodihydrofluorescein diacetate (DCFDA) levels in Control and TSC2KD cells under the conditions and times indicated in the panel. Data points represent n=5 independent experiments plotted as Mean $\pm$ SEM **C.** Representative western blot of compression (10 kPa for 15 minutes) induced eEF2 T56 phosphorylation in control and cells pretreated with 5mM NAC. Ponceau S staining serves as a loading control. Blots are representative of n=2 independent experiments. **D.** Portion of eIF5 sequence showing $CX_2C\text{-}X_{19}\text{-}CX_2C$ motif (top). Alpha-fold generated representation of Zn2+-coordinated cysteine tetrahedral structure found in eIF5 (bottom left). Representative redox-western blot of eIF5 (bottom right). Blots are representative of n=2 independent experiments. **E.** Representative redox-western blot of eEF2. Ponceau S staining serves as a loading control. Blots are representative of n=2 independent experiments.
Paired t-test, two-tailed (B). *P* values: $* P < 0.05$

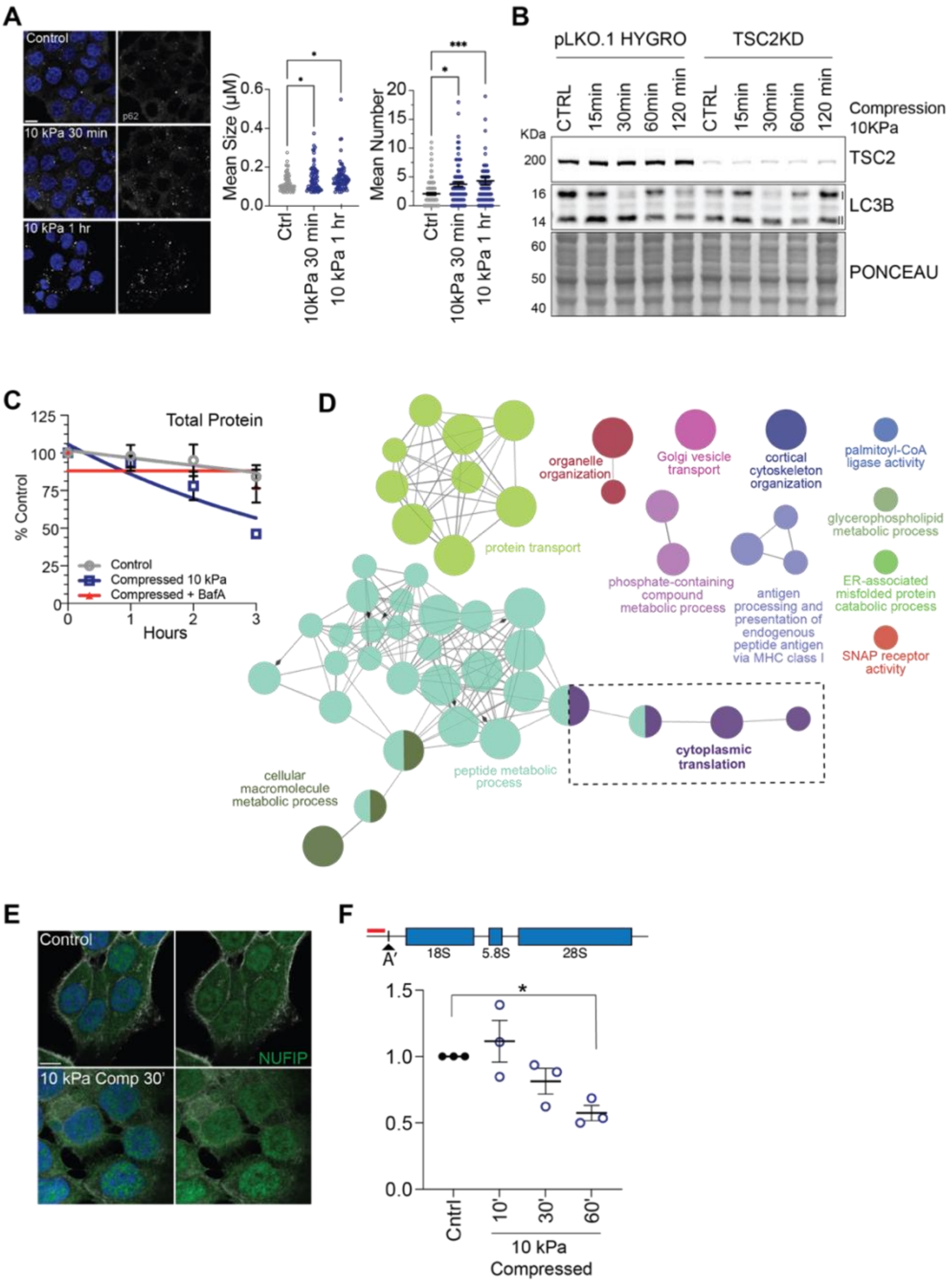


**Supplementary Figure 6. A.** Representative immunofluorescence of p62 foci in control and compressed conditions under compression(left). Scale bar represents 10 $\mu$m. p62 Foci quantification under control and compressed conditions. Both Foci number (right) and size (left) were quantified. Data represents the summed data from three independent replicates which as foci/cell, average size/cell, each dot representing a single cell, respectively: n = 118, 82 and 96 cells for foci number; n = 84, 69 and 80 cells for mean foci area. Data is represented as Mean $\pm$ SEM. (right) **B.** Representative western blot analysis to monitor LC3B lipidation in response to compression in Control and TSC2KD HCT116 cells. Blots are representative of n=2 independent experiments. **C.** Degradation kinetics of L-azidohomoalanine (AHA) labelled proteins in control (grey), compressed (blue) cells, and compressed cells pretreated with 200 nM BafilomycinA (red). **D.** Proteins enriched in TMEM192 immunoprecipitates relative to whole-cell extract (WCE) were analysed using the ClueGO plugin in Cytoscape for GO Biological Process enrichment. The analysis identified enrichment of proteins associated with cytoplasmic translation, supporting the presence of translation-related proteins within the TMEM192-enriched fraction. **E.** Representative confocal imaging of NUFIP1 localization as scored by immunofluorescence for control (Top) and 10kPa compressed (Bottom) conditions. Scale bar represents 10 $\mu$m. **F.** Quantitative Real Time PCR analysis of 47S rRNA. Schematic of oligo(red) placement within the transcript. The oligo was designed to capture non-processed 47SrRNA (Top). Comparison of control and compressed conditions over 0-60 minutes following the application of compression (Bottom). Each point represents an independent replicate. Data is plotted as Mean $\pm$ SEM for n=3 independent experiments.

2-way ANOVA with Sidak's multiple comparison test(B&F). *P* values: $* P < 0.05$, $*** P < 0.001$
